# Epigenomic Profiling and Single-Nucleus-RNA-Seq Reveal *Cis*-Regulatory Elements in Human Retina, Macula and RPE and Non-Coding Genetic Variation

**DOI:** 10.1101/412361

**Authors:** Timothy J. Cherry, Marty G. Yang, David A. Harmin, Peter Tao, Andrew E. Timms, Miriam Bauwens, Rando Allikmets, Evan M. Jones, Rui Chen, Elfride De Baere, Michael E. Greenberg

## Abstract

*Cis*-regulatory elements (CREs) orchestrate the dynamic and diverse transcriptional programs that assemble the human central nervous system (CNS) during development and maintain its function throughout life. Genetic variation within CREs plays a central role in phenotypic variation in complex traits including the risk of developing disease. However, the cellular complexity of the human brain has largely precluded the identification of functional regulatory variation within the human CNS. We took advantage of the retina, a well-characterized region of the CNS with reduced cellular heterogeneity, to establish a roadmap for characterizing regulatory variation in the human CNS. This comprehensive resource of tissue-specific regulatory elements, transcription factor binding, and gene expression programs in three regions of the human visual system (retina, macula, retinal pigment epithelium/choroid) reveals features of regulatory element evolution that shape tissue-specific gene expression programs and defines the regulatory elements with the potential to contribute to mendelian and complex disorders of human vision.

## INTRODUCTION

The human central nervous system (CNS) is comprised of diverse tissues and cell types, whose unique forms and functions arise from their distinct programs of gene expression. The precise timing, levels and cell-type-specificity of gene expression are controlled by cohorts of *cis*-regulatory elements (CREs), including promoters and distal enhancers. These short DNA sequences (100-500 bp) consist of binding sites for sequence-specific transcription factors (TFs) and function coordinately to recruit transcriptional machinery to specific sites in the genome (Nord et al., 2013). Genetic variation within the sequence of CREs, between individuals within a population or between species, can lead to changes in TF binding and thus changes in the expression of linked gene(s). These regulatory variants are now appreciated to play a central role in the evolution of species-specific traits and phenotypic variation between individuals in complex traits including disease (Spielmann and Mundlos, 2016; Zhang and Lupski, 2015) (Gordon and Lyonnet, 2014; Maurano et al., 2012; Villar et al., 2015). A major obstacle toward understanding normal and pathological consequences of regulatory variation in human tissues is that many cell type and tissue-specific CREs have not been comprehensively mapped in the relevant human tissues. Thus, their functions in development and tissue homeostasis remain incompletely understood. Accordingly, the identification and characterization of CREs has become a major focus of genetics in the early 21^st^ century (Consortium et al., 2007).

The extreme diversity of cell types within the human CNS allows for complex sensory processing and behavioral repertoires. The full magnitude of this diversity is still unknown, however large scale efforts to catalog this diversity and determine the cellular basis of CNS diseases are currently underway (Ecker et al., 2017). This diversity poses a fundamental challenge to the study of gene regulation as each cellular phenotype arises from the expression of distinct cohorts of TFs that work together to selectively activate cell type-specific CREs to instruct cell-type specific programs of gene expression. The importance of this problem is underscored by the neurodevelopmental, neurodegenerative and psychiatric disorders that originate from disruptions in gene regulation. Such disorders may arise from mutations in genes encoding TFs, chromatin remodeling factors or mutations affecting CREs themselves (Brandler et al., 2018; Iwase et al., 2017; Short et al., 2018) (Jeong et al., 2008; Son and Crabtree, 2014). Because of the complexity of the CNS, it remains exceedingly difficult to connect regulatory variants to specific phenotypic changes in the molecular, cellular or circuit-level features of the CNS.

The retina is a classic model for CNS development, function, and gene regulation because of its highly stereotyped cellular composition and circuit organization. This reduced cellular diversity compared to other CNS regions has recently made the retina a powerful system for characterizing the mechanisms by which TFs function at CREs to control the specification and differentiation of retinal cell types (Andzelm et al., 2015; Corbo et al., 2010; Hao et al., 2012; Samuel et al., 2014). The retina consists of 6 major cell classes that are specified from a common pool of progenitors during development through the cooperation of broadly expressed and lineage-defining TFs. The majority of cells in the mammalian retina are rod photoreceptor cells, which mediate the earliest steps in vision. The retinal pigment epithelium (RPE) is directly opposed to mammalian photoreceptor cells and is required for photoreceptor cell survival and function. The retina and RPE share an embryonic origin in the early optic vesicle, however as mature tissues they are functionally and molecularly distinct. Together with the macula, a specialized central region of the human retina that mediates high-acuity vision, the retina, macula, and RPE are commonly affected in human visual disorders. It is therefore important to understand the genetic regulation in these tissues to serve as a model of CNS function and disease.

TFs that specify the cell types of the retina and RPE are highly conserved across species. Mutations in TFs that disrupt gene expression in specific cell types often result in similar phenotypes in mice and humans including congenital blindness, photoreceptor degeneration, ocular malformations and glaucoma (Freund et al., 1997; Jordan et al., 1992; Nishimura et al., 1998; RetNet). It is unclear, however, to what extent the CREs that recruit these TF are conserved between species. Indeed, the evolutionary conservation of CREs is significantly lower than that of protein coding genes (Villar et al., 2015). An increasing number of studies show that mutations in specific CREs cause inherited retinal diseases (Bhatia et al., 2013; Ghiasvand et al., 2011; Yang et al., 2006). It is therefore critical to map and characterize the CREs that regulate essential gene expression in human eye tissues to refine the genomic search-space for additional human disease-causing non-coding mutations. These analyses will also facilitate discovery of the mechanisms by which TFs selectively bind and activate cell type-specific CREs to regulate the expression of genes essential for proper retinal development and function.

In this study, we used an integrated epigenomic approach to identify and functionally characterize CREs in three tissue regions that are essential for human vision: the retina, macula and RPE/choroid. By comparing the epigenetic landscape of these tissues, we found that tissue-specific CREs are regulated primarily at the level of differential chromatin accessibility and through distinct cohorts of TFs. We assessed the binding of five TFs that are essential for retinal function to CREs in the human genome and found that rather than being redundant, specific combinations of TFs are essential for CRE function. To investigate the role of TF/CRE interactions in individual retinal cell types, we profiled gene expression in >4,000 single human retinal nuclei and identified cell-type-specific CREs. Comparison of human and mouse retinal CREs demonstrated functional conservation of CREs at essential retinal genes. We also found instances in which the highly dynamic evolution of CRE architecture and TF binding within specific loci led to species-specific gene expression changes in specific retinal cell types. Lastly, by mapping adult and developmental patterns of CREs we were able to show that CRE identification can facilitate the discovery and interpretation of disease-associated regulatory variation in CREs of the human genome. Taken together, this resource identifies and characterizes the CREs responsible for essential gene expression in the adult human visual system. By comprehensively defining the genetic regulatory information necessary for the functioning and maintenance of the human visual system, our data provide new insights into the mechanisms by which regulatory variation shapes the development of complex neural tissues, species-specific traits, and disease outcomes affecting human vision.

## RESULTS

### Identification of active *cis*-regulatory elements in human visual tissues

To identify the specific CREs in the genome that control genes necessary for human visual function we first sought to identify shared and tissue-specific CREs in adult human retina, macula and RPE/choroid based on epigenetic features of enhancers and promoters (Figure 1A&B). Candidate active CREs in each tissue type can be defined by: 1) accessible chromatin as a result of TF binding, 2) enrichment for histone modifications associated with active enhancers or promoters and 3) association with active gene expression (Figure 1B). Chromatin accessibility was determined using the Assay for Transposase Accessible Chromatin (ATAC-Seq) (Buenrostro et al., 2013) on nuclei purified from adult post-mortem unfixed human retina and macula (Figures 1C and S1A). DNAse-Seq datasets were used to identify chromatin accessibility in RPE (Consortium, 2012) (Figure 1C). To distinguish active CREs from other accessible regions, we co-localized accessible regions with the histone mark consistent with active enhancers and promoters, acetylated lysine 27 on histone 3 (H3K27ac), using ChIP-Seq (Creyghton et al., 2010; Rada-Iglesias et al., 2011). Altogether, we identified approximately 30,000, 20,000 and 12,000 regions in the retina, macula and RPE/choroid, respectively (Figure 1C).

**Figure 1.**
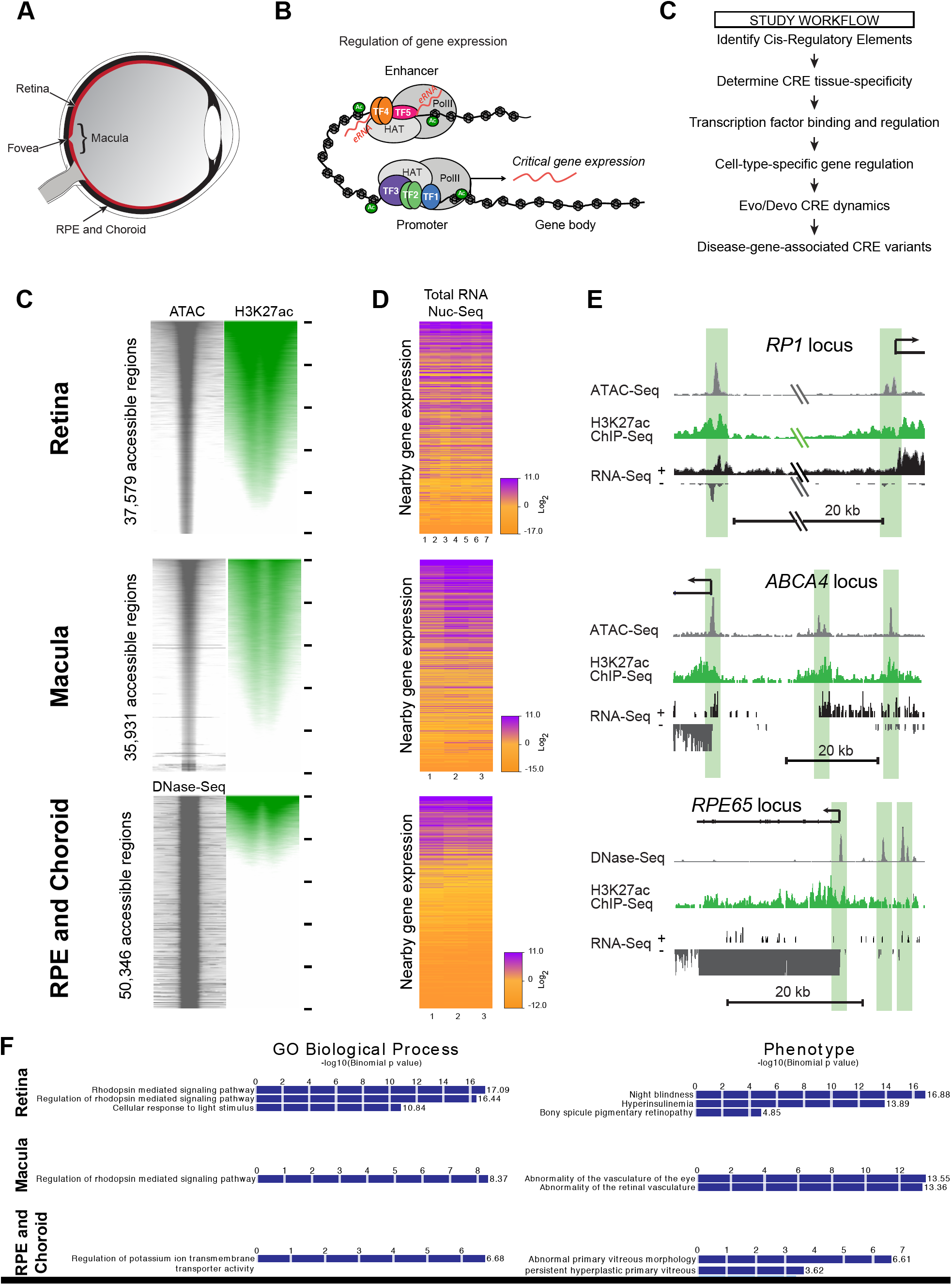
Identification and Characterization of Active Cis-Regulatory Elements in Human Retina, Macula and RPE. **(A)** Schematic of cis-regulatory control of gene expression. Transcription factors (TF1-5) bind in combination to promoters and enhancers to recruit co-factors including histone acetyl transferases (HATs) and basal transcriptional machinery such as RNA polymerase (PolII). HATs acetylate lysine residues on histones, including H3K27ac (Ac). PolII can induce transcription of enhancer RNAs (eRNAs). Both, H3K27ac and eRNAs are associated with active cis-regulatory elements (CREs). **(B)** Schematic cross-section of the human eye with tissues commonly affected in inherited retinal diseases labeled. **(C)** Genome-wide DNA accessibility and H3K27ac in adult human retina, macula and retinal pigment epithelium (RPE) and choroid, by the Assay for Transposase Accessible Chromatin (ATAC), DNase-Seq, and H3K27ac ChIP-seq. Each accessible genomic region (ATAC or DNase-Seq) is represented as a single horizontal line centered on the peak summit, with a window of +/-1kb. ATAC or DNase-Seq signal is plotted in gray. H3K27ac ChIP-Seq signal is plotted in green. For each tissue, windows of DNA accessibility and H3K27ac are ordered on highest to lowest total H3K27ac signal within the 2kb window. **(D)** Expression of genes associated with DNA accessible regions in adult human retina (7 individuals), macula (three individuals) and RPE/choroid (three individuals) as determined by sequencing total RNA from nuclei (Nuc-Seq). Genes are ordered according to their corresponding accessible regions from (C). **(E)** Representative gene loci showing custom UCSC browser tracks for ATAC-Seq or DNase-Seq, H3K27ac ChIP-Seq and total RNA Nuc-Seq from adult human retina, macula or RPE/choroid. **(F)** Enrichment of biological processes and phenotypes associated with candidate active CREs in each tissue according to analysis using the Genome Regions Enrichment of Annotations Tool (GREAT; (McLean et al., 2010)).

Active CREs engage transcriptional machinery to promote target gene expression. To confirm that these identified regions are associated with active gene expression in each tissue, we performed RNA-Seq on adult human retina, macula and RPE/choroid. RNA-Seq was performed on total RNA extracted from cell nuclei (total RNA Nuc-Seq), to avoid cross contamination of RNA between the intercalated cells of the retina and RPE/choroid, and to enrich for enhancer RNAs (eRNAs), a marker of active enhancers (Kim et al., 2010). Total RNA extracted from cell nuclei is also a more direct readout of active transcription than steady state levels of RNA in the cytoplasm. We then compared the level of proximal gene expression to the level of H3K27ac at putative enhancers and promoters. We found that levels of H3K27ac are correlated with levels of proximal gene expression (Figures 1C, 1D, S1D). Highly expressed genes associated with active CREs in each tissue include the known disease genes *RP1, ABCA4* and *RPE65* in retina, macula and RPE/choroid, respectively (Figure 1E). At each of these gene loci, proximal promoters and one or more distal enhancers were identified based on the co-occurrence of DNA accessibility, H3K27ac signal and eRNAs. The correlation of each of these assays was quantified between bioreplicates and across tissues to assess reproducibility and tissue-specificity (Figure S1A, see also Supplemental Information).

Active enhancers and promoters in each tissue contribute to the unique programs of gene expression and biological function of each tissue. To determine the distinct biological functions associated with active enhancers and promoters in each tissue we performed genome regions enrichment of annotations analysis (McLean et al., 2010). We found CREs in the retina and macula to be significantly associated with regulation of rhodopsin signaling and the cellular response to light, consistent with the abundance of photoreceptor cells in both tissues (Figure 1F). Active regulatory elements in the RPE/choroid however were associated with regulation of ion transporter activity, an essential function of RPE to compensate for light-dependent changes in photoreceptor capacitance (Baylor, 1996; Steinberg et al., 1983). This analysis also uncovered association of active CREs with distinct pathologies in each tissue including night blindness, abnormal retinal vasculature, and abnormalities of the vitreous in retina, macula and RPE respectively.

To validate the transcriptional activity of CREs identified using this integrated epigenomic approach, we cloned candidate CREs from the human *ABCA4* Stargardt disease gene locus (Figure S1E) into a luciferase reporter construct and electroporated these constructs individually into the developing mouse retina. The majority of these identified CREs (6/8) increased the transcriptional output of the reporter construct when compared to the empty vector control, suggesting that these elements have intrinsic CRE activity (Figure S1F). Taken together these integrative epigenomic and transcriptomic analyses demonstrate that active regulatory elements contribute to unique biological processes in the human retina, macula and RPE/choroid. Disruption of these regulatory elements by genetic variation could have tissue-specific consequences resulting in visual disorders. The results of these analyses have now defined the sites in the genome in which to look for cis-regulatory variants implicated in disease.

### Tissue-specific gene expression is shaped by DNA accessibility of CREs

The retina, macula and RPE share a common embryonic origin in the nascent optic vesicle, however each tissue performs a distinct function in adult vision. These distinct functions arise from tissue-specific CREs, but how these CREs are determined is unknown. Tissue-specific CREs may be specified when tissue-specific TFs bind to closed chromatin and make it accessible. Alternatively, chromatin may be broadly accessible, but uniquely activated by tissue-specific transcription factors that recruit histone-modifying enzymes and other transcriptional complexes. To distinguish between these mechanisms, we compared CRE accessibility versus CRE activation across the retina, macula and RPE/choroid (Figure 2).

**Figure 2.**
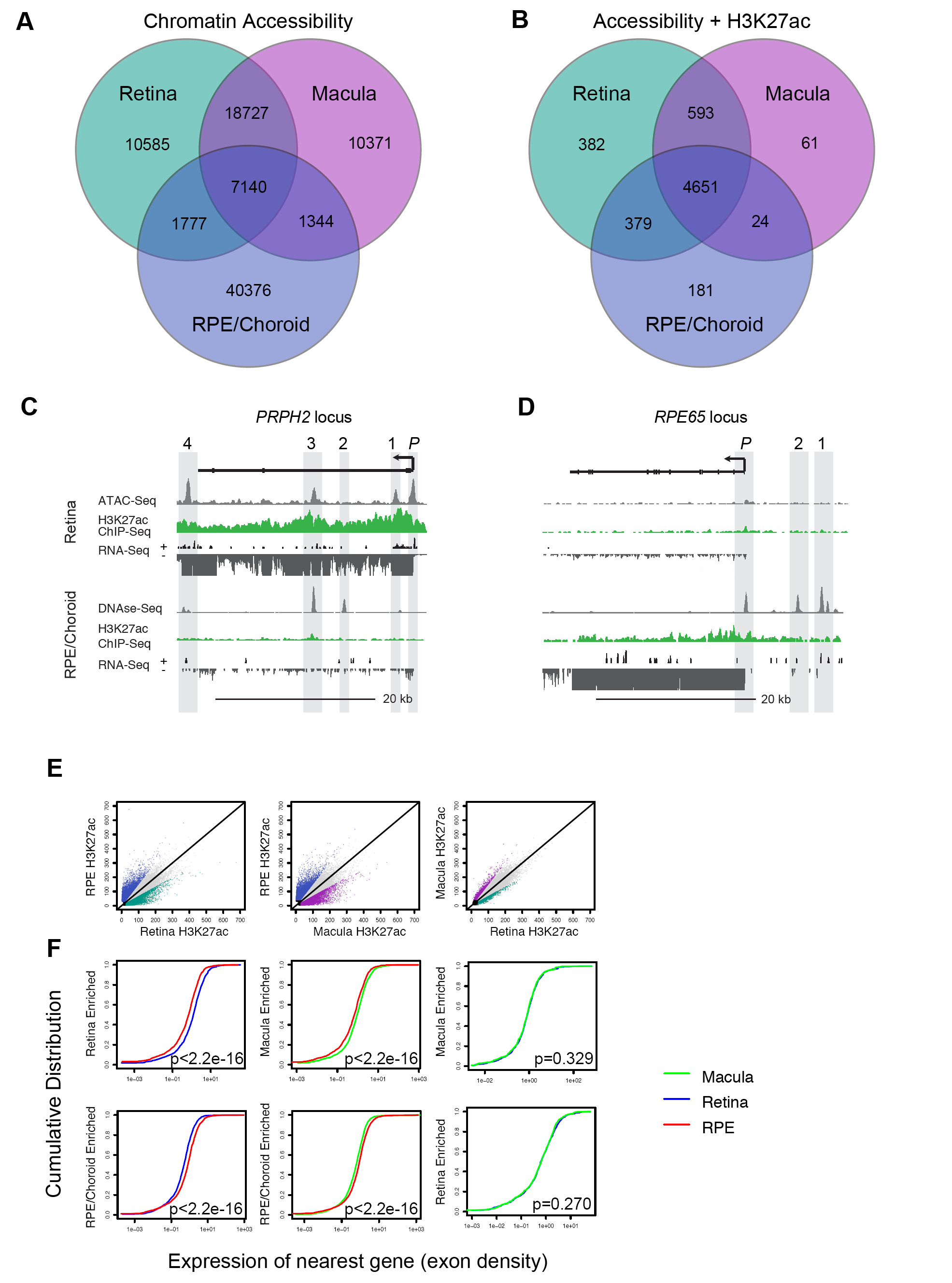
Differential Cis-Regulatory Element Accessibility Drives Unique Patterns of Gene Expression in Human Retina, Macula and RPE. **(A)** Overlap of DNA accessible genomic regions in adult human retina, macula and RPE/choroid. **(B)** Overlap of H3K27ac-enriched genomic regions that share DNA accessibility in adult human retina, macula and RPE/choroid. **(C & D)** Representative tissue-specific gene loci showing custom UCSC browser tracks for ATAC-Seq or DNase-Seq, H3K27ac ChIP-Seq and total RNA Nuc-Seq from adult human retina **(C)** and RPE/choroid **(D)**. **(E)** Pairwise comparison of H3K27ac signal among DNA accessible regions from adult human retina, macula and RPE/choroid (blue: RPE enriched; green: retina enriched; purple: macula enriched). **(F)** Cumulative distribution of gene expression associated with tissue-specific DNA accessible/H3K27ac regions. **(G)** Heatmap of gene expression associated with tissue-specific DNA accessible/H3K27ac regions.

Most CRE accessibility is shared between retina and macula (68% & 69%, reciprocally), likely due to the shared cell types that make up these tissues (Figure 2A). In contrast, only a fraction of their DNA accessible sites is shared with RPE/choroid (23% and 17% reciprocally). This suggests that retina versus RPE/choroid CREs are generally maintained by differential DNA accessibility. Among CREs that are accessible in all tissues, most also share H3K27ac enrichment (74%) (Figure 2B). This shows that while tissue-specific CRE activation does occur, differential accessibility is the dominant pattern of tissue-specific CREs. Examples of both modes of regulation can be observed at the human disease gene loci *PRPH2, RPE65*, and *ABCA4* (Figures 2C & D and S2).

Intriguingly, genes that are expressed in both retina and RPE can also employ tissue-specific CREs giving each tissue “private” control over shared genes. For example we found that ABCA4, the mutated gene in autosomal recessive Stargardt disease (STGD1) was expressed in both human RPE and retina, which to our knowledge has not be previously reported (Figure S2A). One upstream enhancer (1) that is highly active in the retina (Figures S1E and F, S2A, 3D and E) is accessible in RPE, but not enriched for H3K27ac. In contrast two intronic enhancers (7’ and 8’) are accessible and enriched for H3K27ac in RPE/choroid, but not in retina or macula. Independent regulation of disease gene expression by tissue-specific CREs offers one possible explanation for variability seen in human disease phenotypes among individuals with commonly affected genes.

Whether tissue-specific CREs are primarily regulated at the level of accessibility or activation, the end result should be tissue-specific differences in gene expression. To confirm this, we first identified tissue-specific CREs by comparing levels of H3K27ac between each tissue (Figure 2E). Next, we compared the expression of genes associated with these differentially active CREs (Figure 2F). We found that, as a population, genes associated with tissue-specific CREs demonstrated a tissue-specific pattern of expression. These observations confirm the role of differentially active CREs in driving distinct programs of gene expression to impart unique biological functions to each tissue.

### CRE output is determined by combinatorial transcription factor binding

Ultimately, TFs determine the accessibility, activity and function of CREs. Furthermore, genetic variants within CREs that disrupt TF binding motifs may lead to mis-regulated gene expression and disease. It is therefore important to investigate the individual and combinatorial contributions of TFs to CRE function. We first sought to determine which TFs bind to CREs in human retina, macula and RPE/choroid by performing motif enrichment analysis. In the retina and macula we found significant enrichment of OTX2/CRX, MAF/NRL, ROR, and MEF2 motifs, whereas in RPE AP-1, MITF, TEAD and OTX2/CRX motifs were enriched (Figure 3A, data not shown). Many of these TFs have evolutionarily-conserved roles in retinal and/or RPE function and can lead to human visual disorders when mutated (Freund et al., 1997). However, the combinatorial action of these TFs in the human retina is not well understood.

**Figure 3.**
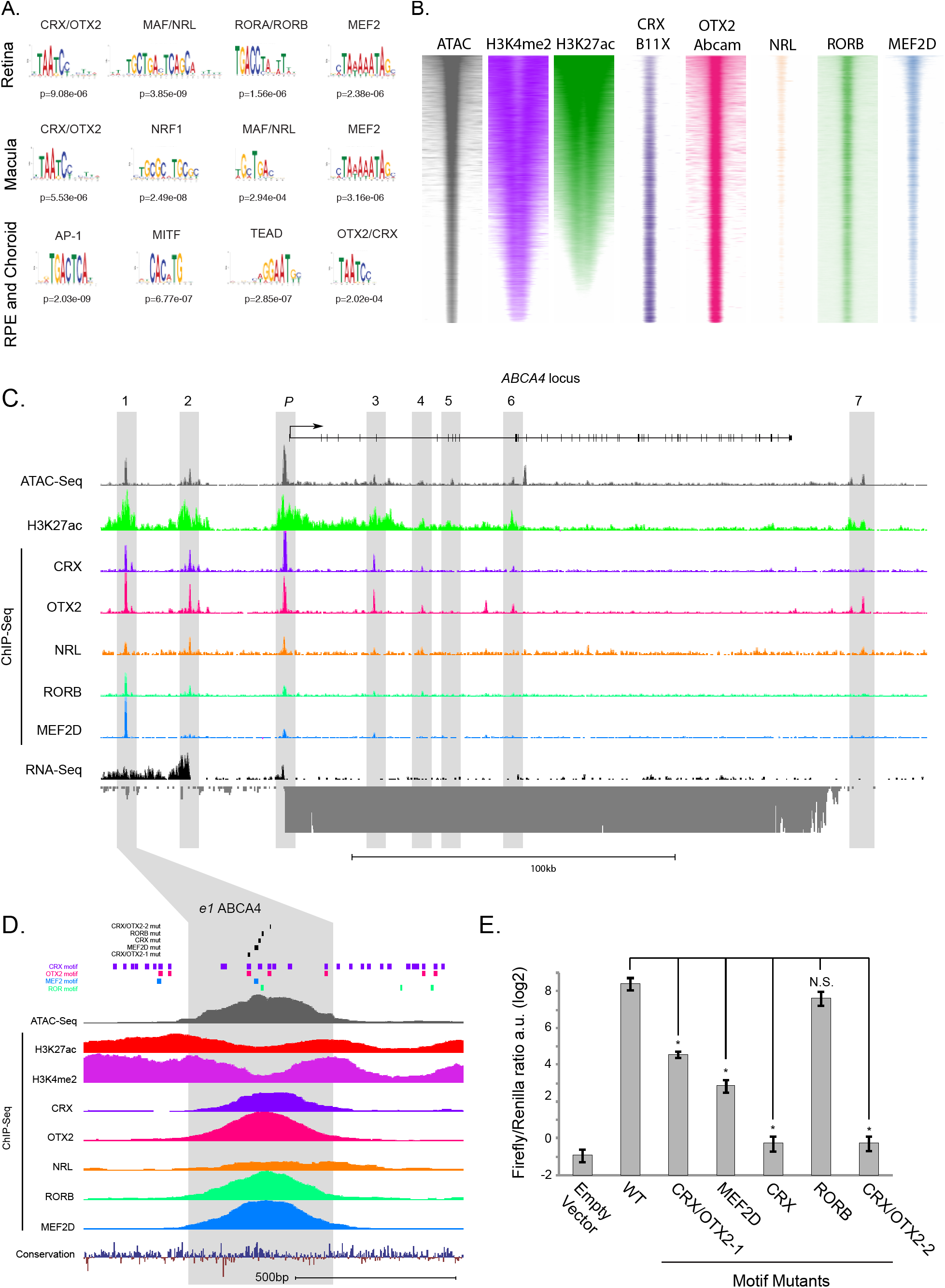
Co-regulation of Human Retinal Enhancers by Combinatorial Binding of Transcription Factors. **(A)** Position weight matrices (PWMs) of TF binding motifs enriched within accessible/H3K27ac+ genomic regions from adult human retina, macula and RPE/choroid. **(B)** Genome-wide distribution of DNA accessibility, H3K4me2, H3K27ac and TF binding in adult human retina by the Assay for Transposase Accessible Chromatin (ATAC) (n=7) and H3K4me2 (n=3), H3K27ac (n=3) or TF ChIP-seq (see methods for n). Each genomic region is represented as a single horizontal line centered on the peak summit, with a window of +/-1kb. Genomic regions are ordered on highest to lowest total H3K27ac signal. **(C)** ABCA4 gene locus showing custom UCSC browser tracks for ATAC-Seq, H3K27ac and TF ChIP-Seq and total RNA Nuc-Seq from adult human retina. Individual candidate CREs (*e1-e7, promoter*) are highlighted in gray. **(D)** A candidate enhancer (*e1*) upstream of the ABCA4 gene showing custom UCSC browser tracks for TF binding motif mutations (black), TF binding motifs (purple: CRX; red: OTX2; blue: MEF2; green: ROR), ATAC-Seq, H3K27ac, H3K4me2 and TF ChIP-Seq and total RNA Nuc-Seq from adult human retina. Area highlighted in gray corresponds to sequence assayed in (E). **(E)** Luciferase reporter assay comparing activity of the consensus human enhancer sequence highlighted in (D) to induced mutations of individual TF binding motifs. (* p<0.05).

Motifs however are only predictive of TF binding, and do not unambiguously identify TFs, as multiple TFs may bind to similar motifs. Therefore, it is important to investigate directly how the binding and activity of TFs relates to human CRE function. We focused on the adult human retina and performed chromatin immunoprecipitation, followed by high-throughput sequencing (ChIP-Seq) for five TFs whose motifs were highly enriched among retinal CREs, CRX, OTX2, NRL, RORB and MEF2D. Antibodies were selected according to previous validation in the literature (Andzelm et al., 2015; Corbo et al., 2010; Hao et al., 2012; Samuel et al., 2014; Wang et al., 2014). Three biological replicates were performed for each antibody unless two validated antibodies were available, in which case two biological replicates were performed per antibody. Anti-RORB antibody was only available in limited amounts and therefore only the top 30% of total peaks were considered for further analysis that matched similar numbers of peaks found genome-wide with the other TFs. In each case, high confidence ChIP-Seq peaks were identified across biological replicates using the ENCODE consortium’s irreproducible discovery rate (IDR) pipeline with a threshold of 1% (Landt et al., 2012). For each of the five assayed TFs, we found that TF protein binding was highly enriched at these active CREs in the adult human retina compared to shoulder regions and genome-wide chromatin input controls (Figure 3B).

To validate the specificity of these antibodies, we performed reciprocal motif analysis for all regions bound by each TF according to ChIP-Seq. In each case, we found that the specific motif corresponding to the antibody-targeted TF was significantly enriched (Figure S3A). Additionally, we found that the paired-type homeodomain TF motif (e.g. CRX, OTX2) was also highly enriched for all TF bound regions suggesting that CRX and/or OTX2 co-bind with these other transcription factors at human retinal CREs. Alternatively, CRX/OTX2 motifs may simply be indiscriminately enriched at sites of binding for any TF in the retina. To distinguish between these possibilities, we performed ChIP-Seq for additional factors CTCF and CREB that are not known to directly regulate photoreceptor cell gene expression. Regions bound by these TFs, were highly enriched for their own motifs, but did not demonstrate enrichment for CRX/OTX2 consensus motifs suggesting that CRX and OTX2 are acting on a specific subset of retinal CREs (Figure S3A). Taken together, these results support the specificity of the antibodies used for these experiments and underscore the combinatorial function of CRX and OTX2 with other TFs in establishing and maintaining photoreceptor enhancers (Andzelm et al., 2015; White et al., 2016).

To examine the relationship of individual TF binding and CRE activity, we ranked human retinal CREs from most active to least active according to H3K27ac enrichment (Figure 3B). We found that the binding of some TFs (OTX2, NRL, MEF2D) was positively correlated with H3K27ac levels suggesting that these TFs or combinations of these TFs may be activating. Oppositely, binding of CRX was highest at the bottom quintile of H3K27ac levels. This suggested that CRX can mediate CRE repression in the mature human retina or is at least bound to inactive sites in the genome (Figures 3B and S3C & D). To further examine these sites, we peformed ChIP-Seq for an additional histone mark H3K4me2, which is enriched at CREs that are either active (H3K27ac-enriched) or poised (H3K27ac-low) (Figure 3B). A repressive role for CRX at CREs is supported by the observation that CRX binds at sites of H3K4me2 enrichment, even when levels of H3K27ac are low (Figure 3B). Furthermore, a repressive role for CRX at some CREs is consistent with a recent study in the mouse retina using massively parallel reporter assays (White et al., 2016). This study, as well as others (Andzelm et al., 2015), did however demonstrate that when CRX binds together with other TFs, the effect can be activating. These observations underscore the significance of combinations of TFs in regulating CREs to promote or repress gene expression.

To assess the significance of specific combinations of TF binding to different aspects of CRE regulation, we determined levels of DNA accessibility, H3K27ac enrichment and nearby gene expression for each permutation of TF binding for five TFs (Figures S3B-D). We found that, as a population, sites where CRX binds without the other assayed TFs have the lowest level of DNA accessibility, the lowest level of H3K27ac and the lowest level of associated gene expression. These sites, however are enriched for H3K4me2, the epigenetic mark of enhancers and promoters (Fig. 3B). This suggests that such sites are maintained in a poised manner, possibly as a vestige of development or as CREs primed to mediate a stimulus-dependent transcriptional response. Among other combinations of TF binding, we did not identify a specific combination that was consistently correlated with levels of accessibility, H3K27ac or nearby gene expression. Taken together this would suggest that regulation of human retinal CREs is heterogeneous and flexible as opposed to fixed according to an invariant order of TF binding. Among models of CRE regulation that have been proposed, these data are more consistent with a flexible “billboard” model of TFs that regulates gene expression rather than an enhanceosome with a fixed spacing and configuration of TF binding and regulation (Spitz and Furlong, 2012).

To further investigate the significance of TF co-binding and to examine the DNA sequence requirements of human CREs, we returned to the *ABCA4* locus. Diverse patterns of TF co-binding were observed across eight highlighted CREs (Figure 3C). Notably, the promoter has the highest ratio of CRX binding to other TFs and also demonstrated low baseline activity by reporter assay (Figure 3C and S1F). In contrast, the upstream enhancer (*e1*) with a lower ratio of CRX binding, but with the highest total observed TF binding (5 out of 5 TFs) was the most highly active CRE at this locus. In order to directly test the contribution of specific TF binding events at this highly active enhancer, we mutated individual TF binding motifs within this enhancer sequence and measured the enhancer activity of these mutants using plasmid-based reporter assays in organotypic retinal explants (Figure 3D).

Determining sequence requirements for CREs is key to interpreting the effect of genetic variants within CREs that may cause disease. Multiple TFs bound to a single CRE may additively or synergistically regulate target gene expression. Alternatively, these TFs may act in a redundant manner to ensure robustness against genetic or cellular perturbations. To distinguish among these possibilities, we mutated individual TF binding motifs at this single, highly active enhancer that binds all five of the TFs that we have examined (Figure 3D). Instead of ensuring robustness, we found that disruption of individual motifs had a profound effect on CRE activity (Figure 3E). Surprisingly, even motifs that occurred multiple times within the same CRE like the shared CRX/OTX2 motif were not redundant. The only motif that did not appear to be required for full CRE function was the RORB motif. These observations are notable as they show that each TF binding event at this highly active enhancer is contributing in a non-redundant manner to enhancer activity, and that CRX appears to function as a transcriptional activator in this context. Taken together, we find that the combination of TFs that bind to individual enhancers can significantly effect CRE activity. This activity may be regulated at the level of DNA accessibility, or CRE activation and can interact with additional CREs to affect target gene expression. These observations are informative for interpreting the impact of genetic variation within human CREs.

### Deconvolution of cell-type-specific regulation

Profiling of epigenomic features from intact tissues of the human CNS is confounded by the extensive cellular heterogeneity of the brain The human retina is less heterogeneous than other CNS regions, however it still contains a considerable diversity of cell types (Masland, 2001). The signal arising from epigenomic assays performed on the intact retina is likely a function of the abundance of each cell type within the tissue and the strength of the signal within that cell type. Nonetheless, cell-type-specific CREs may still be identified by focusing attention on CREs associated with cell-type-specific gene expression (Aldiri et al., 2017). The abundance of different cell types in the human retina and the patterns of cell-type-specific gene expression however, have not been extensively characterized. Single-cell-RNA-sequencing is a powerful new tool to deconvolve cell types in complex tissues (Klein et al., 2015; Macosko et al., 2015). To identify cell-type-specific CREs we first sought to determine the abundance of different cell types in the human retina and to identify cell-type-specific gene expression in human retinal cells. Then with knowledge of the genes that are selectively expressed in a given cell type it should be possible to assign the nearby CREs to that cell type.

We optimized the protocol of Klein and colleagues (Klein et al., 2015) to profile individual cell nuclei (single cell Nuc-Seq) from adult human retinas from three unrelated individuals. By profiling RNA from single nuclei, we avoid transcriptional contamination caused by cell-to-cell contacts and obtain a more accurate snapshot of active transcription. This also obviates the need for intact, living cells because intact nuclei can be efficiently isolated from flash frozen human tissue, making it possible to use stored tissue from biobanks. Using the t-distributed stochastic neighbor embedding (tSNE) algorithm, we identified clusters of cells that based on expression of known marker genes corresponded to all major cell classes of the mature retina including rod and cone photoreceptors, horizontal, bipolar, Müller glial, amacrine and retinal ganglion cells (Figure 4A) (Bumsted O’Brien et al., 2004; Ferda Percin et al., 2000; Gong et al., 2006; John et al., 2000; Satija et al., 2015; Wu et al., 2012). These classes were identified retrospectively based on patterns of known marker gene expression (Figure S4A). Rod photoreceptor cells were the most abundant class of cells profiled, suggesting that the majority of the ATAC-Seq and ChIP-Seq signal originates from epigenomic features in this cell type (Figure 4B). However, the abundance of other cell classes in the human retina was higher than that of non-rod cells observed from single cell analysis of the mouse retina (Macosko et al., 2015). Notably, cone photoreceptor and horizontal cells made up a larger percentage of all cells in the human retina compared to mouse, consistent with classical neuroanatomical studies of rodents and primate and rodent retinas (Jeon et al., 1998). Of note, tSNE identified clusters of cells representing subtypes of different cell classes including different subtypes of bipolar, amacrine and retinal ganglion cells. Diversity within these classes has been well documented in mouse, including in single cell profiling studies from our group and others (Cherry et al., 2009; Trimarchi et al., 2007).

**Figure 4.**
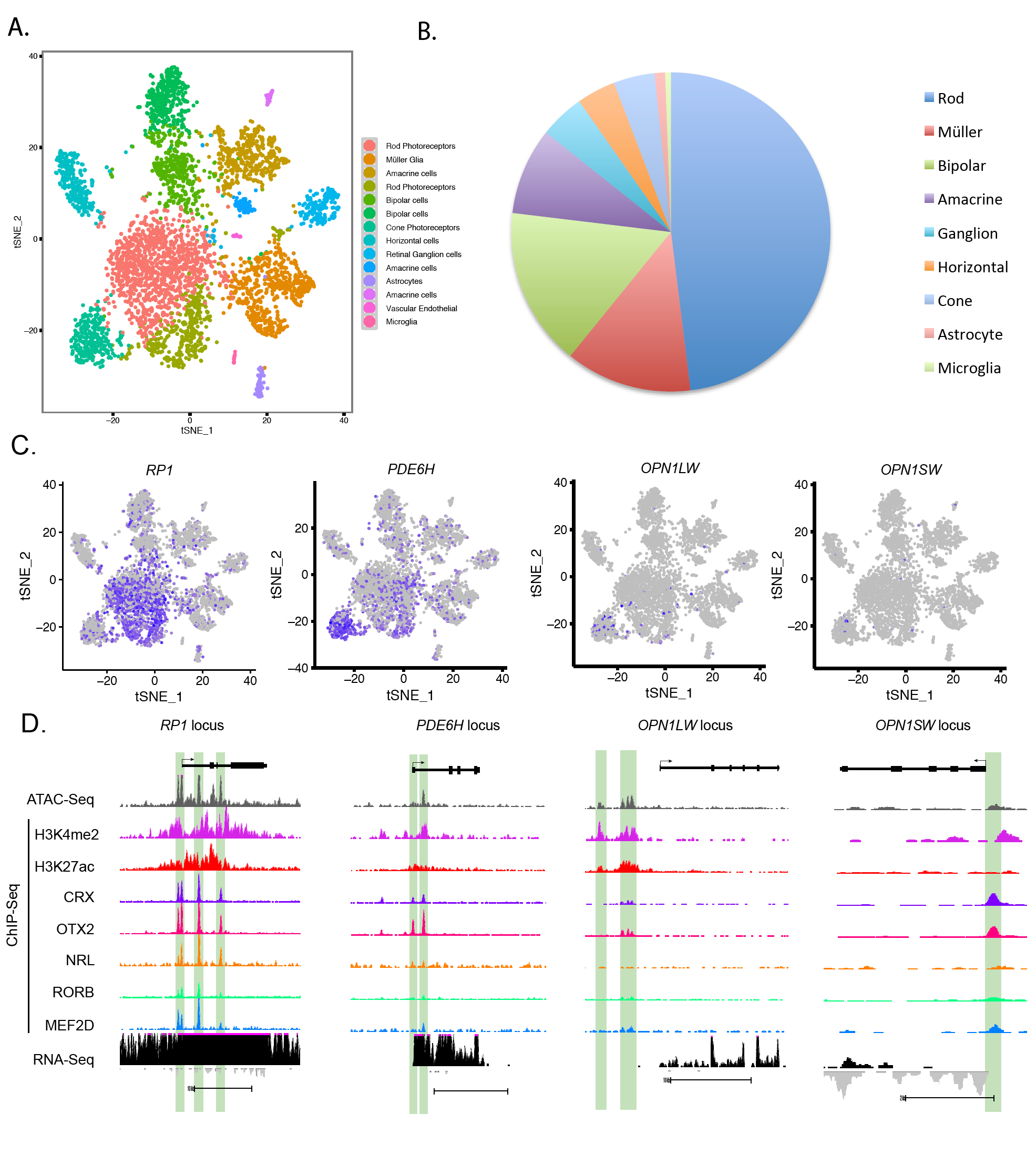
Deconvolution of Cell-Type-Specific Regulatory Elements. **(A)** t-Stocastic Neighbor Embedding (tSNE) plot of global gene expression of 4,763 single nuclei from adult human retinal cell nuclei; nuclei are colored according to 14 unsupervised clusters. **(B)** Proportion of cell classes in the human retina identified according to known marker gene expression in single retinal nuclei (n=3 bioreplicates) (see also Figure S4A). **(C)** Marker gene expression of individual photoreceptor cell types (*RP1*: rod; *PDE6H*: pan-cone; *OPN1LW*: long-wave length [red] cones; *OPN1SW* short-wave length [blue] cones.). **(D)** Epigenetic features at photoreceptor cell type-specific marker gene loci shown by custom UCSC browser tracks for ATAC-Seq, H3K4me2, H3K27ac and TF ChIP-Seq and total RNA Nuc-Seq from adult human retina.

To determine if cell-class-specific CREs could be identified, we verified the expression of cell-class-specific marker genes in our single-Nuc-Seq dataset (Figure S4A). We then examined these individual loci in our dataset to assess DNA accessibility, histone modifications and TF binding (Fig. S4B). We found that the following cell class markers maintained their specificity of enrichment in the human retina and had observable signatures of CREs at their gene loci: *RP1* and *NR2E3* (rod photoreceptors), *ARR3* and *PDE6H* (cones), *ONECUT2* (horizontal cells), *VSX2, PRKCA* and *GRM6* (bipolar cells), *CLU* and *APOE* (Müller glial cells), *GAD1* (amacrine cells), *SCL17A6*/*NEFL* (Ganglion cells), *GFAP* (astrocytes) (Figures 4 and S4 and data not shown). These genes represent markers of broad classes of cells in the retina. To determine if CREs associated with lowly abundant cell types were identifiable in our dataset we analyzed gene loci of cone photoreceptor cell type markers *OPN1LW* and *OPNSW*. Each of these gene loci demonstrated nearby accessible regions. In the case of *OPNSW*, the promoter region was accessible with binding of CRX, OTX2 and MEF2D. At the *OPNLW/MW* region, the well-studied upstream locus control region (LCR) (Carroll et al., 2010) was clearly accessible and binding of CRX, OTX2 and MEF2D was observed. Together these findings suggest that cell class-and cell type-specific regulatory elements can be identified in these integrative datasets although they are generated using heterogeneous retinal tissue, even from cell types that represent less than 7% of the total tissue.

To examine the profile of cell-class-specific gene associated CREs as a population we identified the top 100 most-enriched genes expressed in each cell class whose promoter regions demonstrated accessibility by ATAC-Seq (Figure S4C). We then quantified the ATAC-Seq and H3K4me2 and H3K27ac ChIP-Seq signal at the promoters of these genes (Figure S4D). For each cell-class-enriched cohort, we observed enriched H3K27ac and H3K4me2. For each cell-class-enriched cohort, we observed enriched H3K27ac and H3K4me2. Together, these data suggested that epigenetic features of cell-class-specific promoters can be observed in datasets collected from intact tissue.

### Conservation and evolution of cell type specific regulatory elements

CREs are important substrates of evolutionary change (Villar et al., 2015). However, the architecture of the visual system is remarkably conserved across species with regard to structure, cell types and the function of critical TFs that specify the cellular diversity during development and reinforce cell type gene expression programs in mature retinal cell types (Arendt, 2003; Gehring, 2002). Thus, conservation of regulatory elements associated with critical visual genes may be considerable. However, CREs in other tissues, such as the liver, are highly dynamic during mammalian evolution (Villar et al., 2015). To assess whether conserved retinal architecture is reflected in the genome in the form of conserved CREs, we examined the conservation between human and mouse CREs at critical visual genes. We first compared the number and topography of candidate enhancers at the mouse and human Rhodopsin locus. Rhodopsin is a G-protein coupled receptor, essential for rod cell phototransduction in all mammals. It is also among the most commonly mutated genes in human inherited retinal disorders (RetNet).

We observed remarkable conservation of regulatory element topography between mice and humans despite extensive evolutionary divergence between the two species (Figure 4A). This topographic conservation is evident from the number and relative location of individual CREs at the *RHO* locus. Additionally, the pattern of TF binding to each CRE appears also be conserved with the exception of MEF2D, which appears to be present at CREs at the human, but not mouse *RHO* locus. Despite this topographic conservation, the conservation of DNA sequence is limited to discrete regions within each CRE (Figure S5A). This demonstrates that regulatory conservation can be maintained in absence of extensive DNA sequence conservation.

To assess conservation of regulatory elements at a gene expressed in a less abundant cell type, we analyzed the *CABP5* locus, a calcium binding protein that is an established marker of bipolar cells in the mouse retina (Figure 5C) (Kim et al., 2008). *CABP5* was originally identified as sharing homology to calmodulin and is closely related to *CABP4*, which is associated with inherited retinal disorders (Haeseleer et al., 2000; Zeitz et al., 2006). A proximal promoter and a downstream enhancer are evident in both mouse and human retinas (Figure 4B). The topography of these elements with respect to the *CABP5* coding sequence is conserved, however the binding of TFs is not. Only OTX2 was bound to the mouse promoter and enhancer, while the human promoter and enhancer have acquired binding of CRX, OTX2, NRL, MEF2D and RORB. Among these factors CRX and NRL are enriched in rod photoreceptors, suggesting that this gene may be differentially regulated in the human retina. To determine if the regulation of *CABP5* has been altered between the mouse and human retina, we analyzed our human single cell transcriptome data and compared this to single cell data acquired from the mouse retina (Macosko et al., 2015). In mouse *Cabp5* expression is largely restricted to bipolar cells, in contrast *CABP5* is expressed in bipolar cells and rod photoreceptors in the human retina (Figure 5C). Of note, a 54bp low complexity (CT) insert in the downstream mouse enhancer may prevent the recruitment of other TFs (Figure S5B). Together these observations show that interspecies differences in gene expression can occur through co-evolution of promoters and enhancers to affect not just the activity of a gene, but also its cell-type-specificity.

**Figure 5.**
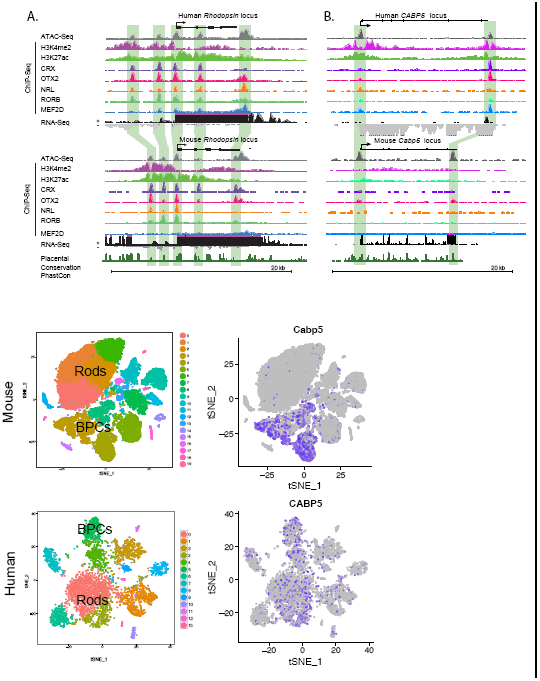
Conservation and Divergence of CRE Function in the Human and Mouse Retina. **(A)** *Rhodopsin* gene locus showing custom UCSC browser tracks for ATAC-Seq, H3K4me2, H3K27ac and TF ChIP-Seq and total RNA Nuc-Seq from adult human or mouse retinas. Individual candidate CREs are highlighted in green. **(B)** *CABP5/Cabp5* gene locus showing custom UCSC browser tracks for ATAC-Seq, H3K4me2, H3K27ac and TF ChIP-Seq and total RNA Nuc-Seq from adult human or mouse retinas. Individual candidate CREs are highlighted in green. **(C&D)** t-Stochastic Neighbor Embedding (tSNE) plot of 31,528 single cells from the mouse retina from (Macosko et al., 2015) (C) or 4,763 single cell nuclei from adult human retinas (D) according to global gene expression; nuclei are colored according to unsupervised clusters (Rods: clusters comprised of rod photoreceptor cells; BPCs, clusters comprised of bipolar cells). **(E-F)** Expression of *Cabp5/CABP5* among single retinal cells/nuclei in the mouse and human retina, respectively. **(G)** *In vivo* expression of TdTomato (red) in the mouse retina driven by the mouse *Cabp5* promoter and enhancer shown in Fig.5B; expression of GFP driven by the *UbC* promoter (GFP), (blue: DAPI). **(H)** *In vivo* expression of TdTomato in the mouse retina driven by the human *CABP5* promoter and enhancer shown in Figure 5B; expression of GFP driven by the *UbC* promoter (GFP), (blue: DAPI).

### Developmentally dynamic CREs underlie distinct biological functions, pathologies and regulatory mechanisms

CREs are highly dynamic during tissue development and regulate discrete processes including proliferation, specification and differentiation (Aldiri et al., 2017; Wilken et al., 2015). The retina is a classic system to study cell type specification and differentiation. To investigate the genetic regulation of these processes in the human retina, we sought to identify developmentally dynamic CREs. Furthermore, visual disorders can arise from gene misregulation during retinal development as well as in the adult retina. It is therefore important to consider developmentally dynamic CREs in conjunction with adult CREs when searching for disease-causing variants.

At 10-12 weeks of human fetal development, the earliest born cell types, the retinal ganglion cells and cone photoreceptors are beginning to differentiate. By 14-17 weeks most neurons of the retina have left the cell cycle and the latest born cell types including rod photoreceptors are differentiating (Hollenberg and Spira, 1973; Spira and Hollenberg, 1973). We compared chromatin accessibility in the adult human retina to that of the developing retina at these two critical stages (Consortium, 2012; Wilken et al., 2015).

Clustering of chromatin accessibility data from these distinct time points revealed three specific patterns that dominated the landscape of developmentally dynamic DNA accessibility (Figure 6A). The majority of sites observed at any time point was shared between the two developmental time windows, but was closed in the adult retina (21,828/44,510: 49%). For example, enhancers upstream of the *ATOH7* gene locus, which is required for the differentiation of early born retinal ganglion cells, exhibited this pattern of chromatin accessibility (Figure 6B). There was also a substantial fraction of CREs that were constitutively open across all time points (16,302/44,510: 36.63%) and CREs that were only accessible in the adult differentiated retina (5,518/44,510: 12.40%), such as those found at the *RHO* locus (Figure 6C). Very few regions were specific to either developmental window (10-12wks: 378 (0.85%) or 14-17wks: 346 (0.78%)) or shared between the late developmental window and adult retina (110 (0.25%)). As expected, almost no sites were accessible in a discontinuous pattern i.e. present at 10-12wks and adult, but absent from 14-17wks. (28 (0.06%)). These patterns suggest highly transient transcriptional programs that are active in human retinal development and demonstrate that CREs continue to be licensed and decommissioned as cells mature.

**Figure 6.**
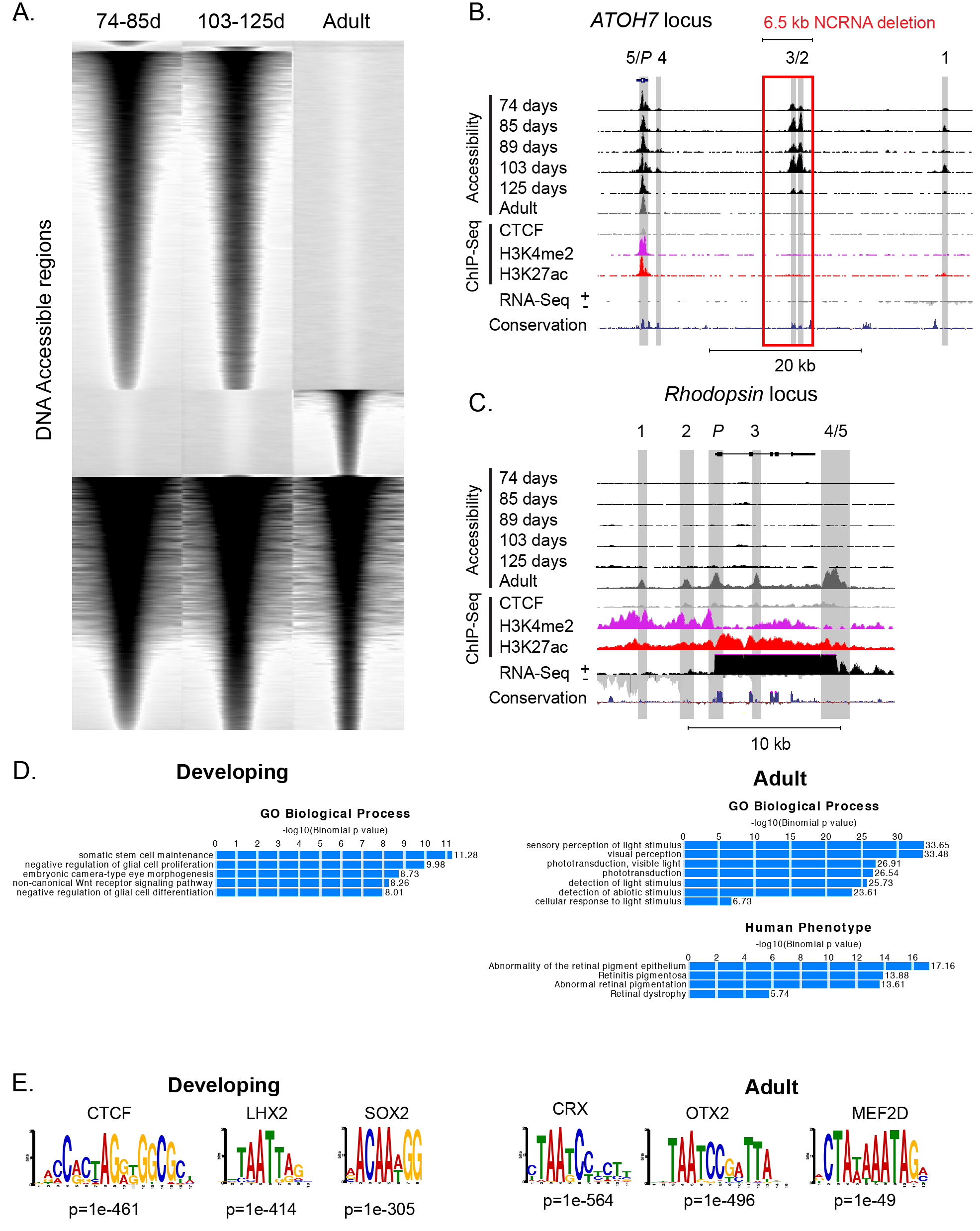
Developmentally Dynamic DNA Accessibility Underlies Distinct Biological Functions, Pathologies and Regulatory Mechanisms. **(A)** Dynamic DNA accessibility during human retinal development. Each accessible genomic region (ATAC or DNase-Seq) is represented as a single horizontal line centered on the peak summit, with a window of +/-1kb. ATAC or DNase-Seq signal is plotted in black. For each developmental stage, windows of DNA accessibility are ordered on highest to lowest total accessibility signal. **(B&C)** Developmentally dynamic DNA accessibility at the ATOH7 or *Rhodopsin* gene loci displayed as custom UCSC browser tracks for DNase-Seq (developing) or ATAC-Seq (adult) from a time course of human retinal development and CTCF, H3K4me2 and H3K27ac ChIP-Seq and total RNA Nuc-Seq from adult human retina (Red Box: 6.5kb deleted ATOH7 enhancer region resulting in inherited human non-syndromic congenital retinal non-attachment; NCRNA; (Ghiasvand et al., 2011)). **(D)** Enrichment of biological processes and phenotypes associated with developmentally dynamic candidate CREs in the human retina according to analysis with the genome regions enrichment of annotations tool (GREAT). **(E)** TF binding motif enrichment at regions that are accessible in developing or adult human retina. **(F)** TF binding at regions of constitutive or adult-specific DNA accessibility.

Identification of dynamic chromatin accessibility is necessary to completely define the genomic search-space for CRE variants that cause disorders of human visual development. To explore this further, we examined developmental chromatin accessibility at two CREs associated with human disease-causing enhancer mutations. Individuals who are homozygous for a deletion of a distal enhancer of *ATOH7* are blind from birth and demonstrate congenital retinal non-attachment (Ghiasvand et al., 2011). Importantly, we found that there are actually two regions of DNA accessibility in the developing human retina encompassed within this deleted region, suggesting that two, not just one, enhancer elements are lost in this disorder (Figure 6B). Additionally, individuals homozygous for a point mutation in a downstream enhancer of *PAX6* demonstrate the congenital eye malformation, aniridia and absence of a fovea (Bhatia et al., 2013). We found that this point mutation also falls within a region of DNA accessibility present during development (Figure S6A). Neither of these enhancers demonstrates accessibility in the adult retina, suggesting that a distinct cohort of genomic regions should be used to search for genetic defects causing developmental disorders rather than disorders that manifest later in life. In contrast, a third disease-associated variant, SNP rs17421627, has been shown to cause developmental retinal vascular abnormalities and is strongly associated with macular telangiectasia type 2 (Madelaine et al., 2018; Scerri et al., 2017). This point mutation falls within an enhancer that is accessible both developmentally and in the adult retina, suggesting how it can contribute to both developmental and later-onset retinopathies (Figure S6B).

The distinct patterns of chromatin accessibility in the developing versus the adult retina further suggest that these elements serve different biological functions and may impinge upon distinct pathologies. We found that regions of accessible chromatin in the developing retina were significantly associated with the biological processes of stem cell maintenance, embryonic camera-type eye morphogenesis and negative regulation of glial development. Adult-specific accessibility, however, was significantly associated with light detection and phototransduction. Apart from DNA accessibility at the *ATOH7* and *PAX6* enhancers, no human disease phenotype terms were associated with regions that are open exclusively in the developing retina. In contrast, adult-specific accessible regions were significantly associated with abnormality of the RPE, retinitis pigmentosa, abnormal retinal pigmentation and retinal dystrophy (Figure 6D).

Distinct patterns of DNA accessibility across retinal development are likely driven by distinct cohorts of TFs. To identify potential regulators of these distinct elements, we looked for known TF DNA binding motifs that were enriched within developmental-or adult-specific accessible regions or within regions that are constitutively accessible. Among the top motifs enriched for developmental-specific accessibility are those that correspond to CTCF, LHX and SOX family TF (Figure 6E). LHX and SOX TF families have well documented roles in the maintenance of progenitor cells during retinal development (Gordon et al., 2013). CTCF is best known for its association with transcriptional insulators and is also the top most enriched motif within the constitutively accessible regions. In the adult-specific accessible regions, the top-most enriched motifs correspond to the transcription factors CRX, OTX2, MEF2D and RORB, which are all known to be essential for proper photoreceptor cell differentiation and function (Figure 6E). Consistent with these analyses, we established that the patterns of TF binding were highly concordant with the motif enrichment and specificity of these distinct regions (Figure S6C). CRX, OTX2, MEF2D and RORB were more frequently bound at adult-specific regions compared to constitutively accessible regions. CTCF in contrast was more robustly bound to constitutively accessible regions compared to regions specifically accessible in the adult retina.

Enrichment of CTCF binding at the large number of constitutively accessible sites suggests that these sites may represent insulator elements. Insulator elements are an important class of CRE that constrain the activity of distal enhancers to a defined set of promoters thus preventing spurious activation of non-target genes (Hnisz et al., 2016). CTCF binding, however, is not specific to insulators as this protein can bind to and act at enhancers, promoters and repressor sites. We therefore used an intersectional approach to identify prospective insulators based on binding of CTCF and exclusion of epigenetic marks for enhancers and promoters (H3K4me2 and H3K27ac). We found that the majority (64%) of putative insulators were constitutively accessible, while only 4% were specifically accessible in the adult human retina.

Taken together, these findings show that unique networks of TFs function at dynamic CREs to regulate stage-specific gene biological functions, whereas the binding of CTCF at insulator elements appears to be largely static and is specified early in human retinal development. Furthermore, these data demonstrate that identified CREs in the human retina overlap with known disease-causing regulatory element mutations. This observation serves as an important proof of concept that mapping CREs can help define the genomic search-space for novel variants that may influence human vision.

### Identification and characterization of functional variants within human cis-regulatory regions

To prioritize the search for novel non-coding variants that affect human vision, we first identified CREs associated with known retinal disease genes (RetNet). Of 253 disease genes, we identified 763 enhancers and promoters (Figure 7A). Genetic variants within these regulatory elements could contribute to disease phenotypes by misregulating the expression of these disease-associated genes.

**Figure 7.**
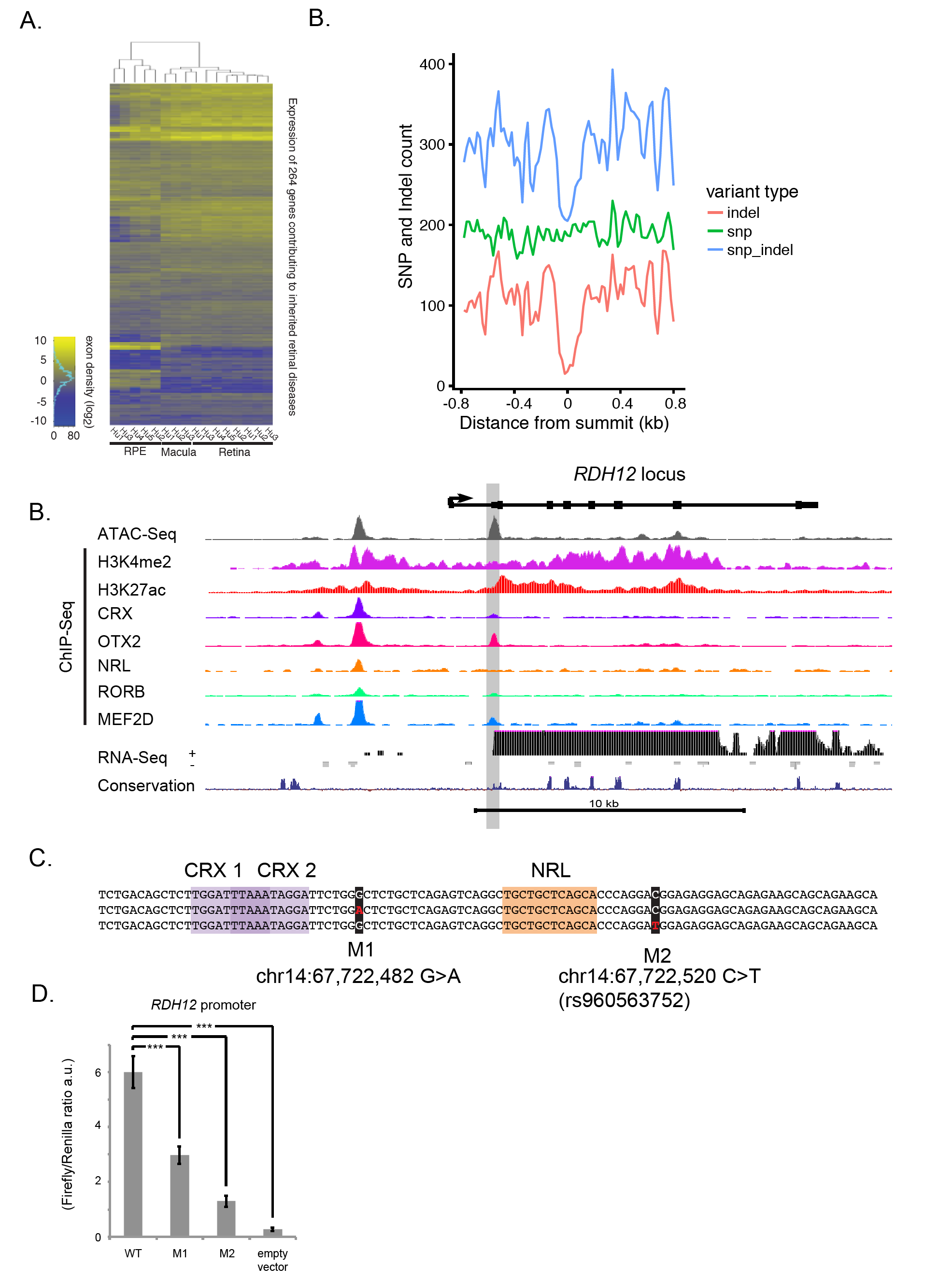
Disease-gene Associated *Cis*-Regulatory Elements and Non-Coding Variation. **(A)** Heatmap of human retinal disease gene expression by tissue type. **(B)** SNP and indel frequency at disease gene-associated CREs (gnomAD, MAF 0.01) (8685 indels, 15306 SNPs). **(C)** The RDH12 gene locus showing custom UCSC browser tracks for ATAC-Seq, H3K4me2, H3K27ac and TF ChIP-Seq and total RNA Nuc-Seq from the adult human retina. Presumptive promoter region highlighted in gray. **(D)** Consensus DNA sequence centered around promoter ATAC-Seq peak summit. Single nucleotide variants (SNVs, red with black highlight) found in unrelated individuals with Leber Congenital Amaurosis/Retinintis Pigmentosa. **(E)** Luciferase reporter assay comparing the relative activities of RDH12 consensus and variant promoter constructs. (*** p<0.0005).

If genetic variants within these CREs are deleterious, then such variants should be rare in unaffected human populations. To assess this, we determined the frequency of SNPs and indels (insertions and deletions) within human retinal-disease gene CREs from approximately 15,500 unrelated individuals without visual disorders (Lek et al., 2016). We found that indels were strongly depleted in the 200bp centered on the peaks of chromatin accessibility compared to flanking regions (Figure 7B). In contrast, no significant depletion of SNPs was observed within these regions. This is consistent with a higher tolerance for single base changes at CREs. Even so, several reports demonstrate that single base changes can have profound affects on enhancer function and disease phenotypes (Bhatia et al., 2013; Madelaine et al., 2018; Scerri et al., 2017).

To identify novel functional variants in human retinal CREs we queried the sequence within these regions from individuals with defined retinal pathologies. These individuals were selected as potentially carrying non-coding regulatory variants because they were found to be heterozygous for a recessive coding pathogenic variant or loss-of-function allele in a known disease gene that matched the disease phenotype. Our hypothesis was that genetic variation within CREs associated with these genes might contribute to the disease phenotype by further lowering the effective dosage of the gene product. For several different unrelated individuals we identified non-coding variants in the CREs of known retinal disease genes including *RDH12, ABCA4* and *MYO7A*.

Two such variants were identified in an active CRE at the *RDH12* locus in individuals with inherited retinal disease (IRD). Biallelic mutations of *RDH12* coding sequence are known to cause recessive Leber congenital amaurosis with severe childhood reitnal dystrophy (Janecke et al., 2004). We found two active CREs at the *RDH12* locus (Figure 7C). The structure of this gene has been predicted to span eight exons with the first exon being non-coding (RefSeq NM_152443). We found no evidence of CRE accessibility or significant DNA sequence conservation at the predicted promoter. Instead we found accessibility at the start of the predicted second exon near the start of the *RDH12* open reading frame. Together with our total RNA-Seq reads, these data suggested an alternative promoter for *RDH12* in the adult human retina. At this candidate promoter we observed local enrichment for H3K27ac and binding of CRX, OTX2, RORB and MEF2D, albeit at moderate levels compared to a nearby upstream enhancer (Figure 7C).

We found two recognizable consensus binding sites for CRX/OTX2 at this prospective promoter. Immediately downstream of these binding sites we found two separate variants in two unrelated individuals, *RDH12 c.-162G>A* (hg38, chr14:67722482G>A) and *RDH12* c.-123C>T (hg38, chr14:67722520C>T) (reported in (Thompson et al., 2005)). Neither were previously characterized as deleterious, however *c.-123C* is conserved across simians and *c.-162A* is conserved down to monotremes. We tested the effect of these variants on the transcriptional activity of this CRE. The consensus CRE demonstrates substantial promoter activity in a plasmid-based reporter assay (Figure 7C & D). However, when each SNP is introduced individually, the activity of this region decreases by ~50% and 80% respectively (p<0.001, n=12, t-test) (Figure 7E).

We tested additional identified variants in the *ABCA4* promoter and an intronic *ABCA4* enhancer and at an intronic *MYO7A* enhancer. Several SNPs assayed at the *ABCA4* promoter had no discernable effect on reporter activity. Surprisingly, however two variants (rs11802887 and rs11806223) that were found together in siblings affected by Stargardt disease demonstrated increased reporter activity individually, with one increase reaching significance (rs11802887) (p<0.05, n=4, t-test) (Figure S7A). In contrast, an *ABCA4* enhancer variant (rs752024867; hg38 chr1:94,079,815T>G) found in an unrelated individual with Stargardt disease showed a significant decrease in reporter activity (p<0.05, n=4, t-test) (Figure S7B). Finally, a distinct SNP found in a *MYO7A* enhancer (chr11:77,172,048 C>T) also demonstrated a significant decrease in enhancer activity (p<0.01, n=4, t-test) (Figure S7C). A common theme among all of these variants is that they do not fall within readily recognizable TF binding motifs, however they do demonstrate conservation across clades. This suggests that despite reduced conservation in CREs compared to coding regions, conservation may be predictive of vulnerable DNA sequences within CREs that contribute to disease phenotypes (Madelaine et al., 2018; Villar et al., 2015).

## DISCUSSION

The majority of human genetic variation falls within non-coding regions of the genome, however these regions remain largely uncharacterized and are not routinely screened for disease-causing mutations. Genome-wide assays to identify CREs based on epigenetic signatures have greatly improved our understanding of functional elements within the non-coding genome, however these assays are difficult to interpret in tissues with a high degree of cellular diversity, such as the CNS. In this study we identify CREs that are active in the adult human retina, macula and RPE/choroid, three tissue regions that are required for vision and that are frequently affected by inherited and acquired visual disorders. The retina is an excellent region within the CNS to demonstrate this approach because of its well-characterized anatomy, cellular composition and relatively limited diversity of constituent cell types compared to other regions of the CNS. We find that many of these CREs demonstrate tissue-specific DNA accessibility and each tissue is enriched for a unique cohort of TF binding motifs. Within the retina, we mapped the binding of five TFs known to be required for photoreceptor function and recognized patterns of TF binding associated with CRE activity. We identified putative cell-type-specific CREs by generating single cell RNA-Seq datasets to compare to our integrative epigenomic data. Furthermore, our studies characterize CREs that harbor non-coding variants that are associated with inherited visual disorders.

Our study focuses on adult human eye tissues and yields new insights into genetic regulation, especially when compared to previous studies in the mouse retina and the developing human retina. (Aldiri et al., 2017; Andzelm et al., 2015; Corbo et al., 2010; Hao et al., 2012; Hartl et al., 2017; Samuel et al., 2014; Wilken et al., 2015). By comparing human and mouse data together, our studies shed light on the evolution of non-coding regulatory elements in the visual system, a remarkably well conserved part of the CNS. By comparing CRE landscapes derived from developmental and adult datasets, it is possible to identify distinct CREs that drive gene expression at each stage in retinal development and to map the binding of TFs that constitute the core transcriptional regulatory circuitry of retinal differentiation. Perhaps most importantly by comparing between developing and adult CREs in the human retina, it is possible to identify and test novel and known genetic variants that are associated with developmental and later onset human visual diseases such as LCA, Stargardt disease and macular telangiectasia type 2.

The datasets generated in the course of these studies can now be used to better understand the normal regulation of genes that are essential for cellular function in this critical CNS structure. One practical application of these data is an improved functional annotation of the genome to aid in the interpretation of non-coding variants found by whole-genome sequencing that may contribute to visual disorders. This will greatly extend upon current candidate gene and exome-sequencing efforts to pinpoint the genetic defects present in patients with retinal disease, especially non-standard presentations that might represent new variants of these well-characterized disorders. Both pursuits would benefit from additional assays such as high-resolution chromatin-conformation capture approaches, which could directly associate distal enhancers with specific target genes. Advances in human retinal organoid culture and genome editing also make it possible to manipulate individual human enhancers to test their contribution to retinal function and pathologies in a targeted and prospective manner.

Taken together our data demonstrate the utility of this integrative genomic approach for the characterization of three complex human tissue regions that facilitate vision. This work fills important gaps in identifying and characterizing the function of non-coding regulatory elements and in providing a biologically relevant framework by which genetic variants can be evaluated and understood.

## EXPERIMENTAL PROCEDURES

Detailed experimental procedures are provided below as supplemental information.

## ACCESSION NUMBERS

GEO accession numbers are pending for data generated in the course of this study. Additional data files that were reanalyzed for this study include the following data from the ENCODE Consortium: human RPE DNase-Seq (ENCBS420ENC, ENCBS893GDP), embryonic human retina (ENCSR820ICX, ENCSR474GZQ, ENCSR621ENC), and 8 week mouse retina (ENCSR000CNW). The following datasets were also reanalyzed for use in Figure 5A&B: OTX2 ChIP-Seq in mouse retina GSE54084 (Samuel et al., 2014), CRX ChIP-Seq in mouse retina GSE20012 (Corbo et al., 2010) and NRL ChIP-Seq in mouse retina www.nei.nih.gov/intramural/nnrldataresource.asp (Hao et al., 2012).

## SUPPLEMENTAL INFORMATION

Supplemental Information includes seven figures and three tables are included with this submission.

## ACKNOWLEDGEMENTS

The authors would like to thank Janine Zieg for assistance with figures, members of the Cherry and Greenberg labs for helpful discussions and suggestions. Sinisa Hrvatin provided in depth assistance with the single-nuc-RNA-sequencing. The authors would also like to thank the Genome Aggregation Database (gnomAD) and the groups that provided exome and genome variant data to this resource. This work was funded in part through the NINDS Javits Award to M.E.G. (R37 NS028829), Lab startup funds to T.J.C., the Ghent University Special Research Fund (BOF15/GOA/011) to E.D.B., the Hercules foundation AUGE/13/023 to E.D.B., the Research Foundation Flanders (G0C6715N and G0A9718N) to E.D.B., the Foundation JED to E.D.B., by Funds for Research in Ophthalmology (FRO) to M.B. The content of this study is solely the responsibility of the authors and does not necessarily represent the official views of the funding sources mentioned.

## AUTHOR CONTRIBUTIONS

Conceptualization: T.J.C., M.Y., and M.E.G. Methodology: T.J.C., M.Y., and M.E.G. Investigation: T.J.C. and M.Y. with technical assistance from P.T. Formal Analysis: T.J.C. and M.Y., with assistance from D.A.H. and A.E.T. Resources: E.D.B., M.B., R.A., R.C., and E.M.J. Writing – Original Draft T.J.C., M.Y., and M.E.G. Writing – Review & Editing: T.J.C., M.Y., M.E.G., E.D.B., M.B., R.C., and E.M.J. Supervision: M.E.G. Funding Acquisition: M.E.G. and T.J.C.

## SUPPORTING FIGURE LEGENDS

### SUPPLEMENTAL INFORMATION

#### MATERIALS AND METHODS

##### CONTACT FOR REAGENT AND RESOURCE SHARING

Further information and requests for resources and reagents should be directed to and will be fulfilled by the Lead Contact, Tim Cherry.

## EXPERIMENTAL MODEL AND SUBJECT DETAILS

### Human Tissue

The Harvard Medical Area Institutional Review Board reviewed and approved our tissue procurement and research protocols (IRB15-1457). All experiements conducted in the course of this study conform to these protocols. Post-mortem human retina, macula, and RPE/choroid from de-identified donors were obtained from Lions VisionGift (Portland, OR, USA). The tissue was provided with de-identified medical records including time and cause of death, post-mortem interval prior to cryopreservation of tissue, age, perceived race and sex. The average post-mortem interval between time of death and tissue collection was 6hrs and the maximum post-mortem interval was 12hrs. The average donor age was 52 (max=60; min=30). None of the donors had a prior medical history of ophthalmological conditions or interventions. Information regarding sex and age accompanies the ATAC-Seq, ChIP-Seq, and RNA-Seq files deposited with the gene expression omnibus (GEO; https://www.ncbi.nlm.nih.gov/geo/).

### Mouse Tissue

The Harvard Medical Area Institutional Animal Care and Use Committee has approved our animal breeding and research protocol (04572). All experiments conform to these protocols. Three to five month old C57BL/6J mice from Jackson Laboratories (Bar Harbor, ME, USA) were used for ChIP-seq, RNA-seq, and ATAC-seq performed in this study unless otherwise noted. The sex of the animals used in each experiment accompanies sequence files deposited with the gene expression omnibus (GEO; https://www.ncbi.nlm.nih.gov/geo/). CD-1 mice from Charles River Laboratories (Wilmington, MA, USA) were used for ex vivo electroporation-based luciferase assays. Animals were group housed in specific pathogen free cages with a 12:12 light:dark cycle and access to food and water *ad libitum* (PicoLab Rodent Diet 5053, LabDiet, St. Lous, MO, USA).

## METHOD DETAILS

### DNA accessibility

#### ATAC-Seq

Flash-frozen tissue was thawed by addition of 700ul of lysis buffer (10 mM Tris-HCl, 10 mM NaCl, 3 mM MgCl2, 0.1% NP-40), homogenized by trituration 10x with a p1000 pipet set to 500ul, dounced in an ice cold RNase-free 2mL glass dounce 10x with a loose pestle and 10x with tight pestle, and transferred back to original 1.5mL LoBind tube (Eppendorf). Dounce was washed with an additional 700ul ice cold lysis buffer and buffer was transferred to tube with sample for a final volume of 1.4mL. Sample was then centrifuged in a microcentrifuge at 4C for 10min at 500 rcf. The pellet was re-eluted in 1.4mL ice cold lysis buffer, transferred to a pre-chilled dounce, homogenized again 12x with tight pestle, and transferred back to 1.5mL LoBind tube. Nuclei were counted on a hemocytometer and the volume of lysis buffer/nuclei suspension needed for 20,000 nuclei was aliquoted into a separate 1.5mL LoBind tube. Samples were centrifuged along with tube containing remaining nuclei (for nuclear localized RNA) at 4C for 5min at 500g (RCF). The supernatant was removed from samples and remaining nuclei. Subsequent ATAC libraries were generated with 20,000 nuclei per sample using the Nextera DNA library prep kit (FC-121-1030; Illumina, San Diego, CA USA) according to the protocol described in (Buenrostro et al., 2013; Buenrostro et al., 2015).

#### DNase I-Seq

Sequence files from the ENCODE Consortium (Consortium, 2012) for human retinal pigment epithelium (ENCBS420ENC, ENCBS893GDP), embryonic human retina (ENCSR820ICX, ENCSR474GZQ, ENCSR621ENC), and 8 week mouse retina (ENCSR000CNW) described in (Wilken et al., 2015) were analyzed in parallel with ATAC-Seq data.

### Chromatin immunoprecipitation

#### Crosslinking

Flash frozen human or mouse tissue was homogenized in crosslinking buffer (10 mM HEPES-NaOH pH 7.5, 100 mM NaCl, 1 mM EDTA, 1 mM EGTA) containing 1% formaldehyde (added immediately before crosslinking) by trituration 10x with a p1000 pipet set to 500ul, dounced in an ice cold RNase-free 2mL glass dounce 10x with a loose pestle and 10x with tight pestle, transferred back to original 1.5mL LoBind tube (Eppendorf), and incubated while rotating gently at room temperature for 10 min. Crosslinks were quenched by addition of glycine to a final concentration of 0.125 M and incubated while rotating gently at room temperature for 5 min. Cells were pelleted by centrifugation 5 min at 1,350 g at 4C. Cell pellets were resuspended once with cold PBS and re-spun for 5 min at 1,350 g at 4C.

#### Nuclei prep

Crosslinked cell pellets were resuspended in cold 5mL L1 buffer (50 mM HEPES-NaOH pH 7.5, 140 mM NaCl, 1 mM EDTA, 1 mM EGTA, 0.25% Triton X-100, 0.5% NP-40, 10% glycerol, protease inhibitors; 10 mM sodium butyrate added for H3K27ac ChIPs) by pipetting and rotated vertically at 4C for 10 min. Pellets were spun for 5 min at 1,350 g at 4C, and the supernatants were aspirated. Pellets were resuspended in cold 5mL L2 by pipetting (10mM Tris-HCl pH 8.0, 200mM NaCl, protease inhibitors; 10 mM sodium butyrate added for H3K27ac ChIPs) and rotated vertically at 4C for 10 min. Pellets were spun for 5 min at 1,350 g at 4C, and the supernatants were aspirated. Pellets were resuspended in cold 1.5 mL LB3 (10mM Tris-HCl pH 8.0, 100mM NaCl, 1mMEDTA, 0.5mMEGTA, 0.1% sodium deoxycholate, 0.5% N-lauroylsarcosine, protease inhibitors; 10mM sodium butyrate added for H3K27ac ChIPs) by pipetting and transferred to polystyrene tubes (Thermo Fisher 352099) for sonication.

#### Sonication

Nuclei pellets were sonicated (Bioruptor Plus, Diagenode) in polystyrene tubes on high power with 36 cycles of 30 sec ‘‘on’’, 45 sec ‘‘off’’. After sonication, Triton X-100 was added to 1% final concentration and sonicated chromatin was centrifuged at 16,000 g for 5 min at 4C. The supernatant was used for preclearing and ChIP. All subsequent steps were performed using DNA LoBind tubes (Eppendorf).

#### Preclearing and antibody-bead coupling

Protein A Dynabeads (Life Technologies) were washed twice with blocking buffer (0.1% BSA in LB3 + 1% Triton X-100) and aliquoted for preclearing and antibody-bead coupling. Antibodies were coupled to beads in 1.8 mL of blocking buffer by vertical rotation at 4C for 4 hr. In parallel, each sample of sonicated chromatin was incubated with an equivalent volume of washed bead slurry for preclearing. Each ChIP was performed in 1.8 mL LB3 + 1% Triton X-100 and rotated vertically at 4C for 16 hr. For input samples, 1% of sample was saved for de-crosslinking and DNA purification.

#### Washes and elution

For each wash, beads were rotated vertically in wash buffer at 4C for 5 min. Beads were washed twice with low salt wash buffer (0.1% SDS, 1% Triton X-100, 2 mM EDTA, 20 mM Tris-HCl pH 8.0, 150 mM NaCl), twice with high salt wash buffer (0.1% SDS, 1% Triton X-100, 2 mM EDTA, 20 mM Tris-HCl pH 8.0, 500 mM NaCl), twice with lithium chloride wash buffer (0.25 M LiCl, 1% NP-40, 0.5% sodium deoxycholate, 1 mM EDTA, 10 mM Tris-HCl pH 8.0), and once with TE (50 mM Tris, 10 mM EDTA). Beads were then incubated at 65C in 200 mL of TE + 1% SDS per sample for 30 min with vortexing every 10 min. Eluted protein-DNA complexes were separated from the beads and incubated at 65C for 16 hr to reverse crosslinks.

#### Purification of immunoprecipitated DNA

Elutions were incubated with 10mg RNase A for 1 hr at 37C, followed by 140 mg proteinase K for 2-3 hr at 55C with shaking. DNA was extracted with 1 volume of 25:24:1 phenol-chloroform-isoamyl alcohol and purified with a Qiagen PCR purification kit. ChIP DNA concentrations were determined by Qubit dsDNA HS Assay Kit (Thermo Fisher).

### Gene expression

#### Total RNA Nuc-seq

To determine RNA expression in adult human retina, macula and RPE nuclei, RNA was extracted from excess nuclei (~80M) from the ATAC nuclear preparation protocol using Trizol (ThermoFisher; 15596026). Prior to RNA extraction, an equal amount of ERCC spike-in control RNA (Mix 2) was added to each sample, following the manufacturer’s specifications. RNA was extracted using the Qiagen RNeasy Mini Kit with on-column DNase I digestion (Qiagen 79254).

#### Single Nucleus RNA-seq

Generation of single-nucleus suspensions were prepared by isolating nuclei from flash frozen human retina as described above and purified using an iodixanol gradient (Optiprep; Sigma-Aldrich, St. Louis, MO, USA) with RNase-inhibitor. After gradient centrifugation, the nuclei were washed in dissociation solution (Hrvatin et al., 2018; Klein et al., 2015) containing 0.04% BSA and were resuspended in dissociation solution containing 0.04% BSA and 15% Optiprep for single-nucleus RNA sequencing using a modified inDrops protocol (Klein et al., 2015).

### Library preparation and sequencing

#### ATAC-seq

ATAC-Seq libraries were prepared as described above and sequenced on the Illumina NextSeq 500 platform with 75 bp single-end reads, or 2 x 150 bp paired-end reads.

#### ChIP-seq

2-40 ng of each ChIP sample was used to prepare libraries with the NuGEN Ovation Ultralow Library System v2 kit per manufacturer’s protocol. Libraries were sequenced on the Illumina NextSeq 500 platform with 75 bp single-end reads, or 2 x 150 bp paired-end reads.

#### Total RNA Nuc-seq

RNA-seq libraries were prepared using the NEBNext Ultra Directional RNA Library Prep Kit (E7420L, NEB) and either NEB rRNA depletion kit (E6310, NEB) or Qiagen GeneRead rRNA Depletion Nano kit (180211, Qiagen). Spike-in RNA (Fisher 4456740) were used for sample-to-sample normalization. Samples were then sequenced on the Illumina NextSeq 500 platform with 2x 150 bp paired-end reads.

#### Single-nucleus RNA-seq

Single-nucleus RNA sequencing (inDrops): One or two libraries of approximately 3,000 cells were collected from each donor retina. inDrops was performed as previously described (Klein et al., 2015; Zilionis et al., 2017), generating indexed libraries that were then pooled and sequenced on the NextSeq 500 (Illumina) platform.

### Luciferase assays & CRE variant analysis

See Figures S1F, 3E, 7D and S7. Standard assays were performed as previously described (Andzelm et al., 2015). Briefly, ~450bp genomic sequence DNA fragments corresponding to identified and variant CREs were directly synthesized (IDT; Coralville, IA, USA) and inserted into pGL4 vectors (Promega; Madison, WI, USA) by isothermal assembly using NEBuilder HiFi DNA assembly master mix according the manufacturer’s instructions (E2621, NEB; Ipswitch, MA, USA). The pGL4.10 reporter backbone was used to test the activity of promoter elements cloned directly upstream of the minimal promoter. The pGL4.23 reporter backbone was used to test the activity of enhancer elements cloned into the SalI/BamHI sites. The pGL4.74 reporter plasmid was used as a coelectroporation control. Postnatal day 0 (p0) CD-1 mouse retinas from pups of both sexes were dissected and electroporated with the reporter plasmids, cultured *ex vivo*, and harvested at day in vitro 11 (DIV11). The explant electroporation protocol was adapted from (Matsuda and Cepko, 2008). Retinas were then washed briefly in ice cold 1x PBS and homogenized in 500μl passive lysis buffer with trituration. Homogenate was snap frozen to promote cell lysis and subsequently thawed for analysis of luciferase activity. Luciferase assays were performed using the dual-luciferase reporter assay system (E1910, Promega; Madison WI, USA) on a BioTek Synergy multi-modal plate reader. CRE sequences are listed in Supplemental Table 1.

## QUANTIFICATION AND STATISTICAL ANALYSIS

### Sequence read data processing (ATAC-Seq, ChIP-Seq)

#### Read processing and alignment

Demultiplexed FASTQ files were trimmed with Trimmomatic (version 0.33) using the parameter SLIDINGWINDOW:5:30 (Bolger et al., 2014). Trimmed reads were indexed and aligned to the human or mouse genomes (Human GRCh38/hg38 assembly December 2013; Mouse GRCm38/mm10 assembly, December 2011) using the Burrows-Wheeler Aligner (bwa) tool (Li and Durbin, 2010) with the parameters: bwa aln -q 0 -t 4 -n 2 -k 2 -l 32 -e -1 -o 0 and bwa samse -n 5. Tag directories of reads were created using HOMER (version 4.6) makeTagDirectory (Heinz et al., 2010).

#### Peak calling

High confidence peaks across concordant biological replicates were identified via ENCODE’s Irreproducible Discovery Rate (IDR) pipeline using a 1% IDR threshold (Landt et al., 2012). For each set of replicates, bam files of reads were pooled and 2 pseudoreplicate files were generated by samtools view with the parameters -h -b -s 1.5 and -h -b -2.5. Peaks in biological replicate, pooled replicate, and pseudoreplicate samples were called over pooled input samples using MACS2 (version 2.1.1) (Zhang et al., 2008) with the parameters –nomodel -g <hs or mm> -p 1e-1 –extsize 200, where hs or mm refers to human or mouse genomes. An optimal final set of IDR-filtered peaks was obtained from the peak caller and used as the final high-confidence peak set for further analyses and the centers of these peaks were used as summits for analyses with HOMER.

### Sequence read data processing (Total RNA Nuc-Seq)

#### Read processing and alignment

Reads were trimmed to 70bp and aligned to the human genome (Human GRCh38/hg38 assembly December 2013) using the Burrows-Wheeler Aligner (bwa) tool. Two sets of target sequences were provided and incorporated into the bwa index in addition to the usual 23 chromosomal targets: (1) the human mitochondrial genome (GenBank accession NC_012920.1); and (2) a set of ~10 million short (≤138bp) exon-exon splice-junction sequences (see below). For RNA-seq data, a third set of 92 short (< 2.1kb) spike-in oligos representing a wide range of reference concentrations (ERCC RNA Spike-In Mix, Life Technologies; Mix 2) was also incorporated into the index. Typically ~90% of all reads were mappable, allowing up to 2 mismatches, and of these ~70-85% were mapped uniquely. Multiple reads whose 5□ ends were assigned to the same locus were not flattened to a single count.

The splice-junction target sequences were based on the NCBI RefSeq database for GRCh38. For each annotated transcript, we noted all subsets of two or more exons, not necessarily adjacent, that could be spliced together to produce a sequence at least as long as the read length. Each of these sequences were then trimmed to the maximum number of bases such that a read mapping to the sequence would necessarily cross these ordered exons’ splice junction(s). This procedure produced a library of all unique sets of exons whose intragenic splice junctions could possibly be covered by a read of the given length, based on the RefSeq annotation of exonic loci. Aligned reads thus had the opportunity to align either to genomic (chromosomal) sequences or to exon-junction-crossing sequences found only in mature mRNA.

#### Expression level quantification

An in-house software tool, MAPtoFeatures (Gray et al., 2014), was used to quantify expression levels for individual genes as follows. A database of genic features (CDSs and UTRs) was constructed from ~103,000 genomic transcripts annotated in RefSeq for GRCh38. Merged genes were constructed by unioning all exons in all transcripts assigned to each distinct gene; the resulting segments defined the gene’s exonic coordinates used here (with the gaps between them defining introns). Genes with zero CDS exons were labeled ‘‘noncoding’’. These 24,644 genes were supplemented with 37 mitochondrial genes and 1,752 additional noncoding genes specified by the loci of all ribosomal RNA genes obtained from RepeatMasker (where the options Variations and Repeats, rmsk.repFamily=‘‘rRNA’’ yielded 414 LSU-rRNA_Hsa; 74 SSU-rRNA_Hsa;,1264 5S). The purpose of this step was to allow the filtering out of reads stemming from transcription of repeats and rRNA genes, which tend to get populated to inconsistent degrees from sample to sample depending on the variable quality of rRNA depletion. Reads that aligned uniquely were then queried for their intersection with the exonic ranges of any of the above 26,433 genes, including exon-exon splice junctions. The total number of read bases that overlapped an exonic range was divided by the range’s length to give an average exonic read density (i.e., coverage). All reads were assigned to genes or to intergenic regions. However, only those reads not assigned to noncoding genes counted towards the total normalization count N, which ultimately afforded a more stable comparison of expression levels between samples than simply using the total number of reads. All read densities were normalized to a reference total of 10 million reads and a reference read length of 35 bp through multiplication by the factor (10^7^/N)*(35 bp/70 bp). (Division of these normalized densities by 0.35 yields expression levels in alternative units of reads per kilobase of transcript per million mapped reads, RPKM.) For downstream analyses, genes with a normalized exon density value of 0 in any replicate at any timepoint were discarded to remove lowly expressed genes.

### Sequence read data processing (Single-nucleus RNA-Seq)

#### inDrops sequencing-data processing

Transcripts were processed according to a previously published pipeline (Hrvatin et al., 2018; Klein et al., 2015). Briefly, this pipeline was used to build a custom transcriptome from the Ensembl GRCh38 genome and annotation with Bowtie 1.1.1 (Langmead et al., 2009), after filtering the annotation gtf file (gencode.v17. annotation.gtf filtered for feature_type=‘gene’, gene_type=‘protein_coding’ and gene_status=‘KNOWN’). Read quality control and mapping against this transcriptome were then performed. Finally, unique molecular identifiers were used to reference sequence reads back to individual captured molecules, thus yielding values denoted UMIFM counts. All steps of the pipeline were run with default parameters unless explicitly specified.

#### Quality control for cell inclusion

Our initial dataset contained ~30,000 cells with more than 1,000 reads assigned to each cell. All mitochondrially encoded genes were removed from the dataset. Cells with fewer than 400 or more than 15,000 unique-molecular-identifier counts were next excluded, thus yielding 4,763 high quality cells isolated from three donors.

#### Dimensionality reduction and clustering

All 4,763 cells were combined into a single dataset. The Seurat R package (Macosko et al., 2015; Satija et al., 2015) was used to cluster cells. The data were log normalized and scaled to 10,000 transcripts per cell. Variable genes were identified with the MeanVarPlot() function, which calculates the average expression and dispersion for each gene, then bins genes and calculates a z score for dispersion within each bin. The following parameters were used to set the minimum and maximum average expression and the minimum dispersion: x.low.cutoff=0.0125, x.high.cutoff=3, y.cutoff=0.5. Next, PCA was carried out, and the top 20 principal components were retained. The clustering resolution was set to 0.6. This method resulted in 14 initial clusters across 4,763 cells.

### Quantifying bioreplicate correlations

To validate the reproducibility of our epigenomic and transcriptomic assays, we compared the correlation of bioreplicates performed on distinct samples for ATAC-Seq/DNase-Seq, H3K27ac ChIP-Seq and total RNA Nuc-Seq experiments. For each assay, we found a high degree of correlation between samples of the same tissue type (Figure S1A). As expected, retinal and macular samples were also highly correlated with one another. As anticipated, RPE samples were distinct in every assay, because RPE does not share the same cell types that retina and macula do. Another source of differences in DNA accessibility in retina and macula versus RPE may come from intrinsic differences between ATAC-Seq and DNase-Seq protocols. To evaluate this, we performed ATAC-Seq in mouse retina and compared with existing DNase-Seq data from ENCODE. We found that these datasets generated using different techniques were well correlated (r=0.69, Spearman) and that the profiles of accessibility at representative gene loci were nearly identical (Figures S1B and C). A notable difference however was that while the width of ATAC-Seq windows is highly correlated with H3K27ac signal, DNase-Seq regions appear to be less correlated (Figure 1C).

### Characterization of CREs

#### Identification of active CREs

Active CREs were defined for the purposes of this study as regions of overlapping ATAC-seq or DNase-seq peaks with H3K27ac-ChIP-seq peaks using BEDtools intersectBed (Quinlan and Hall, 2010), using the –wa option to maintain the coordinates from either the ATAC-seq or DNase-seq data. These findings are summarized in Supplemental Table 2.

#### Tissue-Region-Specific CREs

See also Figures 2A&B. BEDtools intersectBed (Quinlan and Hall, 2010) wasg used to identify the extent of overlap in regions of chromatin accessibility and the extent of overlap between regions of H3K27ac-enrichment among shared accessibility regions.

#### Cell-type-specific promoters

See also Figure S4C. Cell-type-specific promoters were identified first by analyzing cell-type-specific gene expression in our single-nucleus RNA-seq data using the differential expression function of Monocle2 (Qiu et al., 2017). The promoter regions of the top 100-most enriched genes for each cell type were identified in our ATAC-Seq data and peak summits were used to center aggregate plots for ATAC-Seq and H3K4me2-and H3K27ac-ChIP-Seq read counts as described in Andzelm, Cherry, et al. (2015).

#### CRE annotation and analysis

See also Figures 1F and 6D. Annotation of of CREs was performed using HOMER annotatepeaks.pl (Heinz et al., 2010) and the Genomic Regions Enrichment of Annotations Tool (GREAT) (McLean et al., 2010) under default parameters. Prior to CRE annotation with GREAT we transposed hg38 bed coordinates to hg19 using the UCSC liftOver tool (https://genome.ucsc.edu/cgi-bin/hgLiftOver) (Hinrichs et al., 2006).

#### Disease-gene associated CREs

See also Figure 7B. Active CREs were associated with specific genes using GREAT(McLean et al., 2010) and HOMER annotatepeaks (Heinz et al., 2010). CREs associated with genes implicated in retinal disease as of 2017 (RetNet) were defined as disease-gene associated CREs for the purpose of this study. These findings are summarized in Supplemental Table 3.

#### Motif enrichment analysis

See also Figures 3A, 6E, and S3A. HOMER findMotifsGenome.pl (Heinz et al., 2010) was used to query known and *de novo* position weight matrices (pwms) for enrichment within the central 200bp of each CRE class as defined by ATAC-seq using parameters we have previously described (Andzelm, Cherry, et al. 2015).

### Conservation analysis

See also Figure S5. 150bp windows were selected within human retina ATAC-Seq peaks to encompass the region of highest local conservation across human, chimpanzee, rhesus, mouse, rat, dog, and opossum. Motifs were identified based on their assigned pwm from the following references (Berger et al., 2008; Harding and Lazar, 1993; Kuo and Calame, 2004; Rehemtulla et al., 1996).

### Developmental accessibility analysis

DNase-seq data downloaded as fastq files from the ENCODE project (Consortium, 2012) was processed and aligned and peaks were called on these data using the MACS2 workflow with IDR as described above. Bed files for early embryonic time points (74 & 85 days) were concatenated together, as were bed files for later embryonic time points (103 & 125 days). Read densities in 10bp bins across 2kb windows centered at summits of interest were obtained for early, late and adult peaks using HOMER annotatePeaks.pl with the parameters hg38 -size 2000 -hist 10 (tag directories were normalized to 10 million reads by default). Differential peaks were identified as being 3-fold increased in the category of interest, with a maximum read density of 50 in the alternate category.

### Data visualization

#### Fixed line plots

See also Figures 1C&D, 3B, 6A & 7A. Tag directories of concordant biological replicates were merged using HOMER makeTagDirectory to create a single tag directory of pooled reads. Read densities in 25bp bins across 2kb windows centered at summits of interest were obtained using HOMER annotatePeaks.pl with the parameters hg38 -size 2000 -hist 25 –ghist (tag directories were normalized to 10 million reads by default). Resulting data matrices were then plotted using JavaTreeView (Saldanha, 2004).

#### Heatmaps

See also Figure S1A. Correlation heatmaps for DNA accessibility or H3K27ac were constructed as follows: High confidence ATAC-Seq or DNase-Seq peaks *from retina, macula, or RPE* were pooled together, defining a list of 123,856 genomic targets for quantifying *DNA accessibility and H3K27ac enrichment*. Every peak’s range was reduced to a standard width of 1,001 bp centered on the ATAC/DNase-Seq peak summit. ATAC-Seq was performed on 8 bioreps of human retina, 3 of macula, and data from 2 bioreps for DNase-Seq of RPE was downloaded from the ENCODE Consortium. H3K27ac ChIP-Seq was performed on 8 samples: 3 bioreps of human retina, 3 of macula, and 2 of RPE. These sequenced reads were trimmed to 70bp and aligned to the human genome (Human GRCh38/hg38 assembly, December 2013) using bwa *as described above*, allowing up to *2* mismatches. For each sample, all uniquely mapped reads from both strands were intersected with the set of ATAC/DNase-Seq target ranges, the compilation of which yielded a relative enrichment level (RPKM) for each range in each sample. Overall renormalization per sample was not necessary, as such factors do not affect correlations between samples. For each pair of samples, the Spearman correlation coefficient was calculated over all ranges having a nonzero expression level in both samples. The resulting correlation values *r*_S_ ranged from 0.576 to 0.948 and are represented in the heatmap of Figure S1A by a color scale covering the range *r*_S_ = [0.5,1.0].

Correlation heatmaps for RNA-seq (Figure S1A) were constructed as follows: RNA-seq was performed on 7 retina, 3 macula, and 3 RPE bioreplicates. Sequenced paired-end reads were trimmed to 70 bp and aligned to the human genome (hg38) using bwa as described above, allowing up to 2 mismatches. Only reads that mapped uniquely, and those for fragment end #1, were retained in subsequent analyses. Reads were intersected with 26,433 genomic targets comprising RefSeq-annotated merged genes, each of which was obtained by unioning its transcripts’ exons, mitochondrial genes, and ribosomal RNA genes, plus a library of short exon-splicing sequences, whose expression contributed to their parent gene’s exonic expression level. In each sample target genes were filtered out if they had a normalized exon density less than 0.20 or a read count (rounded-up sum of fractions of any reads overlapping any exons) less than 3; rRNA expression was also removed from this calculation; the remaining 15,502 genes had informative expression levels in one or more samples. For each pair of samples, the Spearman correlation coefficient was calculated over all genes having a nonzero expression level in both samples. The resulting correlation values *r*_S_ ranged from 0.464 to 0.961 and are represented in the heatmap of by a color scale covering the range *r*_S_ = [0.4,1.0].

#### Genome broswer tracks

See also Figures 1E, 2C&D, 3C, 4D, 5A&B, 6B&C, 7B. Bed files from processed and aligned sequence reads were extended to 200bp and normalized to 10M reads using BEDtools (version 2.23.0) (Quinlan and Hall, 2010) genomeCoverageBED using the -scale parameter before being converted into bigWig format for display on the UCSC genome browser (https://genome.ucsc.edu/).

#### Aggregate plots

See also Figures S1D, S4C, S6C. Tag directories of concordant biological replicates were merged using HOMER makeTagDirectory to create a single tag directory of pooled reads. Read densities in 10bp bins across 2kb windows centered at summits of interest were obtained using HOMER annotatePeaks.pl with the parameters hg38 -size 2000 -hist 10 (tag directories were normalized to 10 million reads by default).

#### Cumulative distribution plots

See also Figures 2F and S1D. For cumulative plots involving RNA-seq data (see Figure 2F), CREs were assigned to target genes by HOMER annotatepeaks.pl. Corresponding exon density values are rank normalized and ordered for plotting cumultative distribution with R. To define upper and lower quintiles, normalized tag count densities from HOMER annotatepeaks are used.

#### Scatterplots

See also Figures 2E and S1B. Tag directories of normalized read counts were averaged across biological replicates. Scatterplots display log_2_ normalized values of these averages unless otherwise noted.

## DATA AND SOFTWARE AVAILABILITY

All sequencing data generated for this study has been deposited into GEO: accession numbers pending. All custom software developed or used including in this study is available upon request.

### KEY RESOURCES TABLE

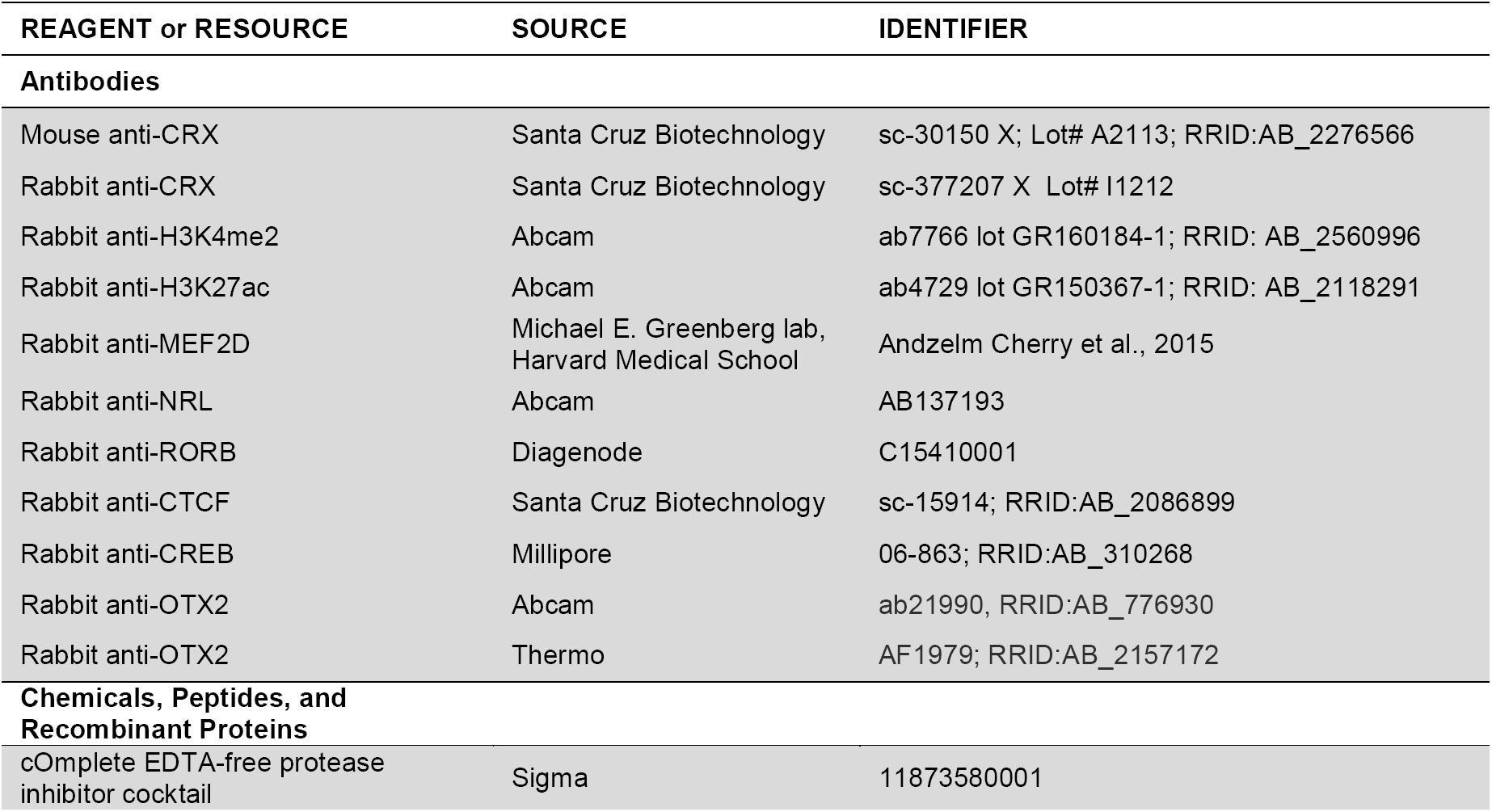
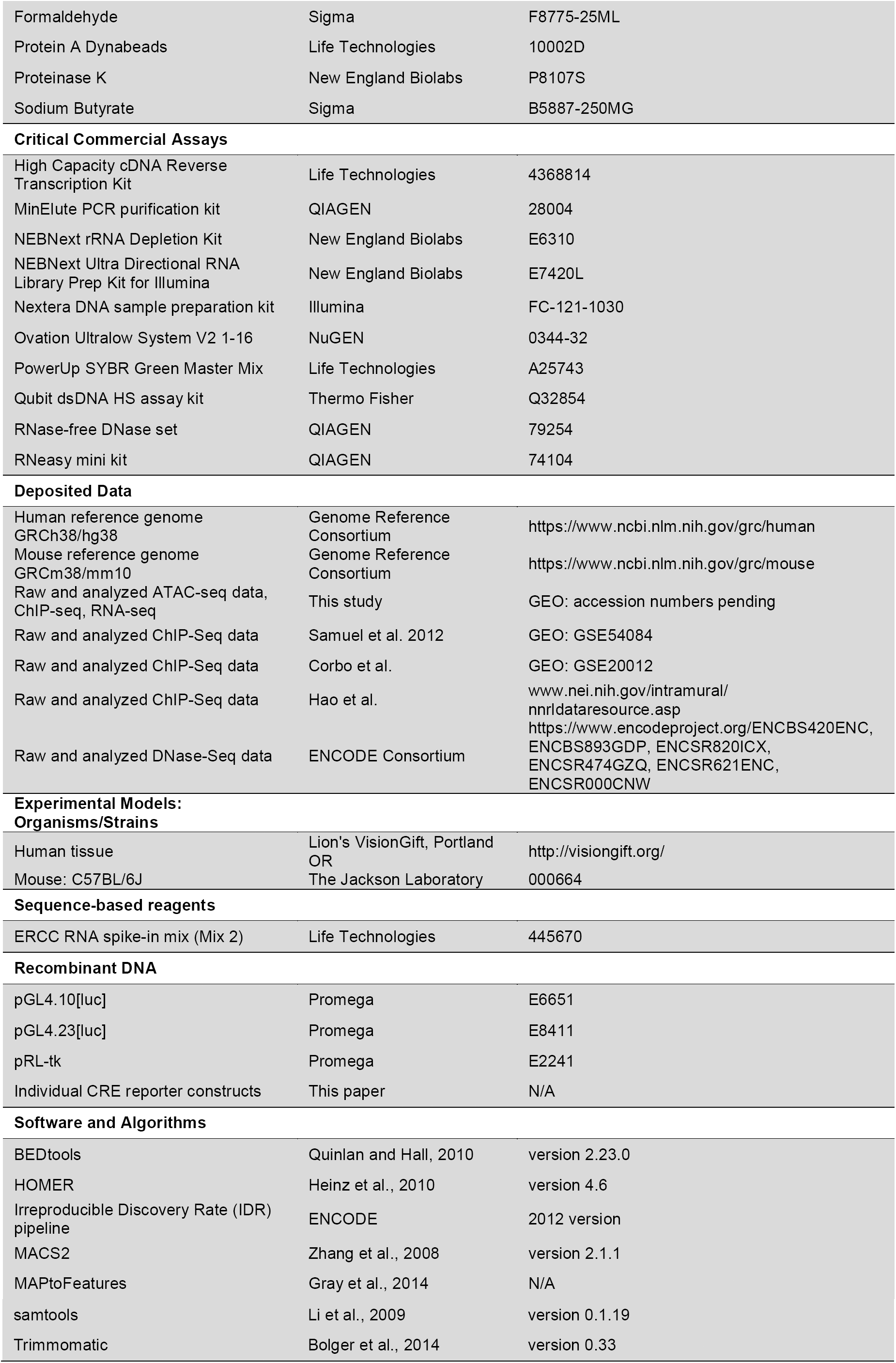

## SUPPLEMENTAL TABLES

**Supplemental Table 1. Luciferase reporter construct sequences.**

**Supplemental Table 2. Proposed Active Cis-Regulatory Elements from Retina, Macula, and RPE/Choroid Hg38 coordinates.**

**Supplemental Table 3. Proposed Disease-Gene-Associated Active Cis-Regulatory Elements Hg38 coordinates.**

**Figure S1.**
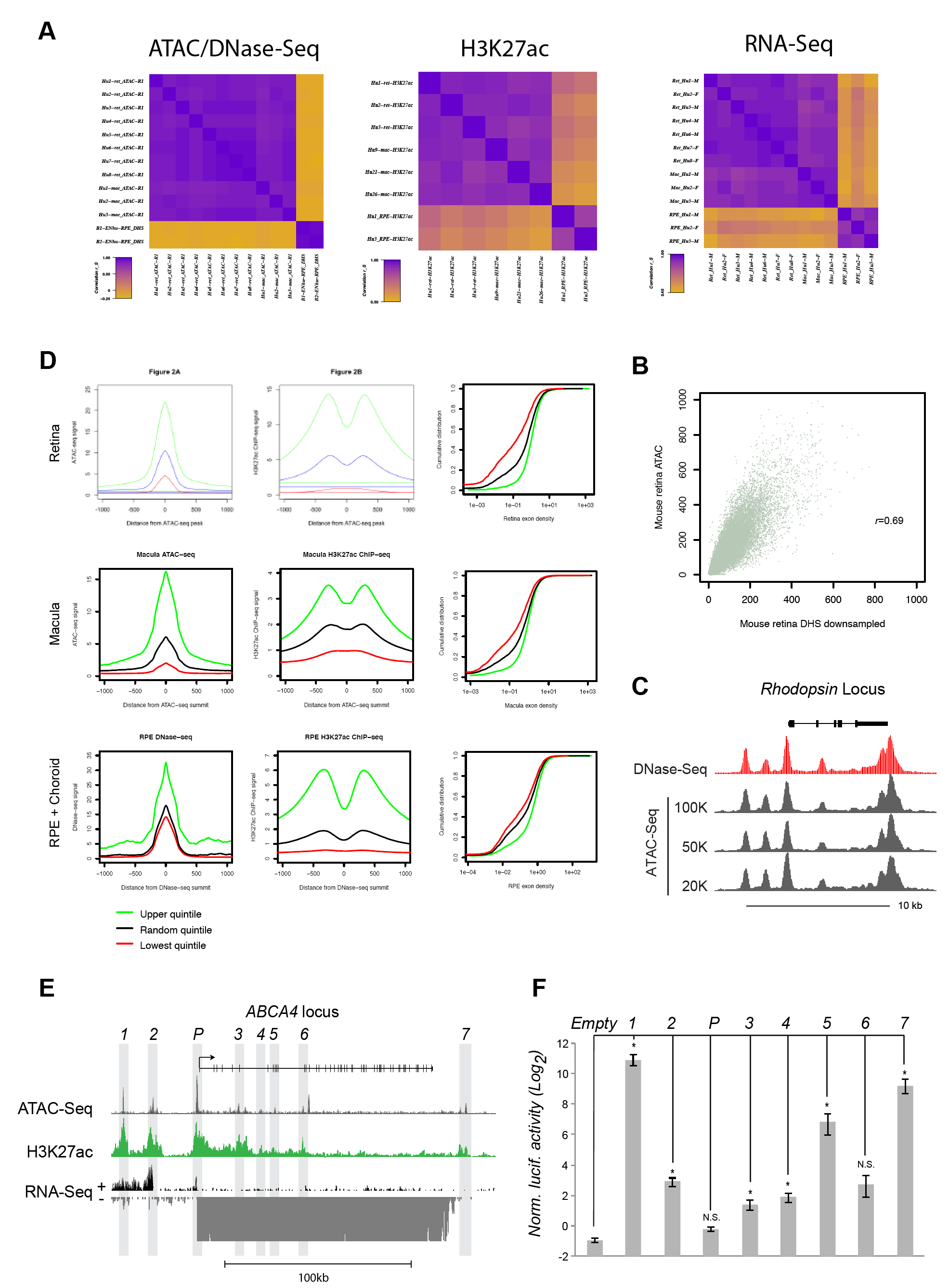
Related to Figure 1. **(A)** Heatmaps of Spearman correlation of ATAC-Seq/DHS, H3K27ac ChIP-Seq and total nuclear RNA-Seq from retina, macula and RPE. **(B)** Correlation plot of ATAC-Seq vs. DHS-Seq from adult mouse retina. **(C)** The *Rhodopsin* gene locus showing custom UCSC browser tracks for DNAse-Seq and ATAC-Seq. **(D)** Aggregate plots of the top (green), bottom (red) and randomly selected (black) quintiles from Figure 1C; Cumulative distribution plots of RNA-Seq of genes associated with top (green), bottom (red) and randomly selected (black) quintiles of H3K27ac at CREs from Figure 1C&D. **(E)** *ABCA4* gene locus showing custom UCSC browser tracks for ATAC-Seq, H3K27ac and total RNA Nuc-Seq from adult human retina. Individual candidate CREs (*e1-e7, promoter*) are highlighted in gray. Luciferase reporter assay comparing the relative activities of identified CREs at the *ABCA4* gene locus to an empty reporter construct. (* p<0.05, n=4).

**Figure S2.**
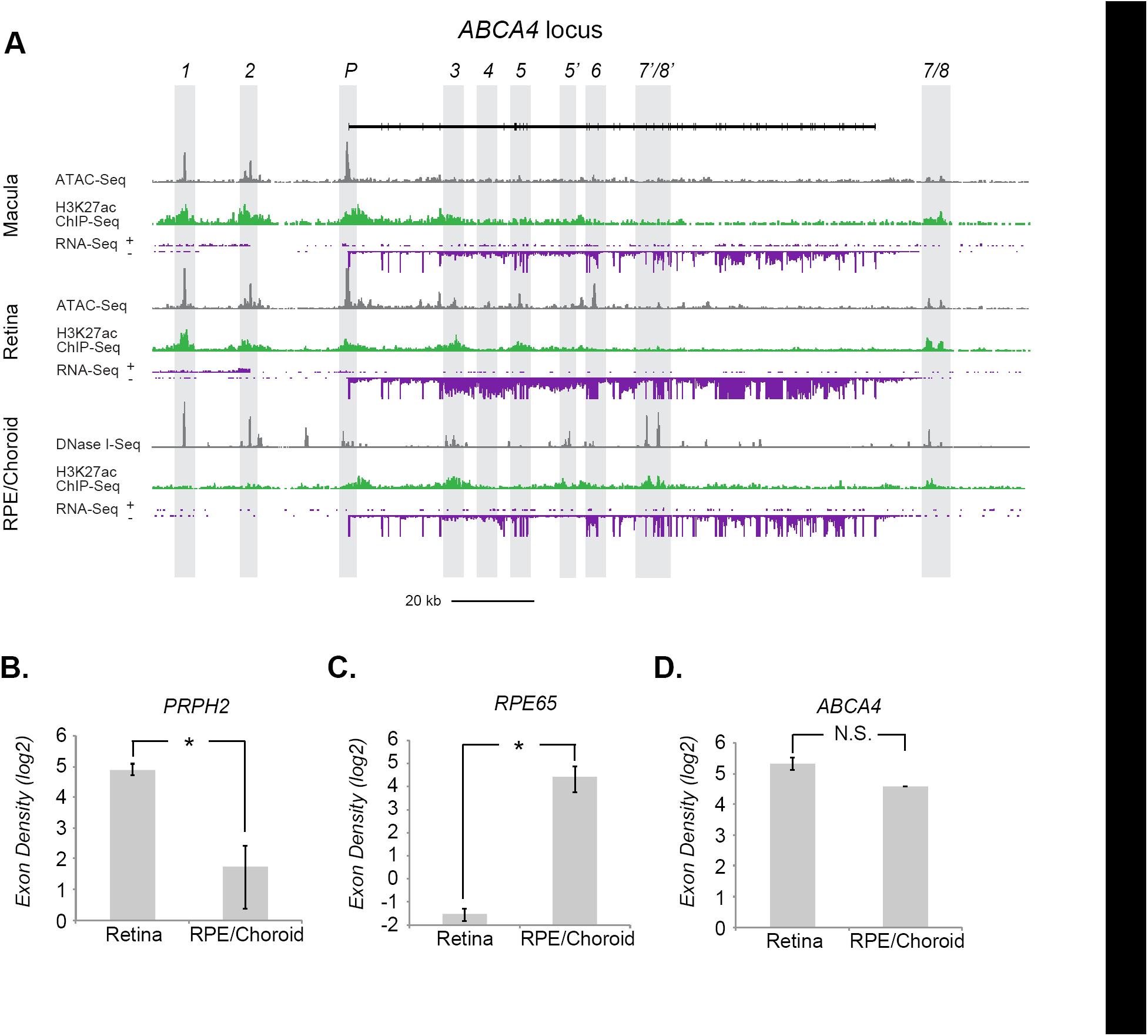
Related to Figure 2. **(A)** *ABCA4* gene locus showing custom UCSC browser tracks for ATAC-Seq, H3K27ac and total RNA Nuc-Seq from adult human retina, macula and RPE/choroid. Individual candidate CREs (*e1-e7, e5’-e7’, promoter*) are highlighted in gray. **(B-D)** Average exon density and standard error of *PRPH2, RPE65* and *ABCA4* expression in adult human retina (n=7) and adult human RPE/choroid (* p<0.05; n=3; t-test, error bars: S.E.M.).

**Figure S3.**
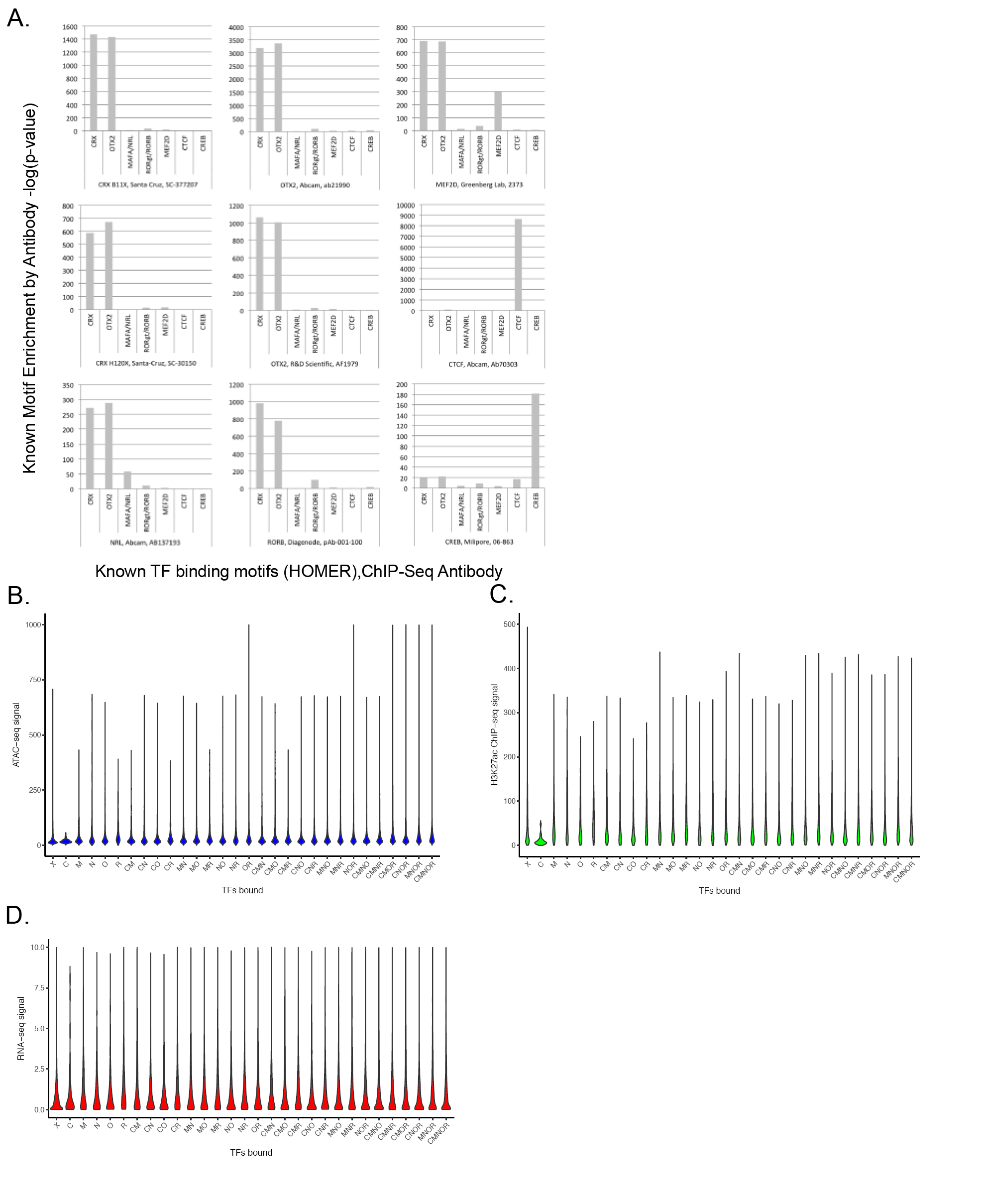
Related to Figure 3. **(A)** Position weight matrix (PWM) of TF binding motifs enriched within ATAC+/H3Kme2-/H3K27ac genomic regions from adult human retina, macula and RPE/choroid. **(B)** Aggregate plot of CTCF binding at ATAC+/H3Kme2-/H3K27ac genomic regions. **(C)** Comparative enrichment of 7 TF binding motifs (CRX, OTX2, MAFA/NRL, RORg/RORB, MEF2D, CTCF and CREB) within ChIP-Seq positive regions identified with CRX, OTX2, NRL, RORB, MEF2D, CTCF and CREB antibodies. **(D&E)** Comparative ATAC-Seq or H3K27ac read density (reads/peak/bp) at sites of combinatorial TF binding. **(F)** Comparative RNA-Seq rank-normalized exon density of genes associated with sites of combinatorial TF binding (C: CRX; O: OTX2; N: NRL; R: RORB; M: MEF2D).

**Figure S4.**
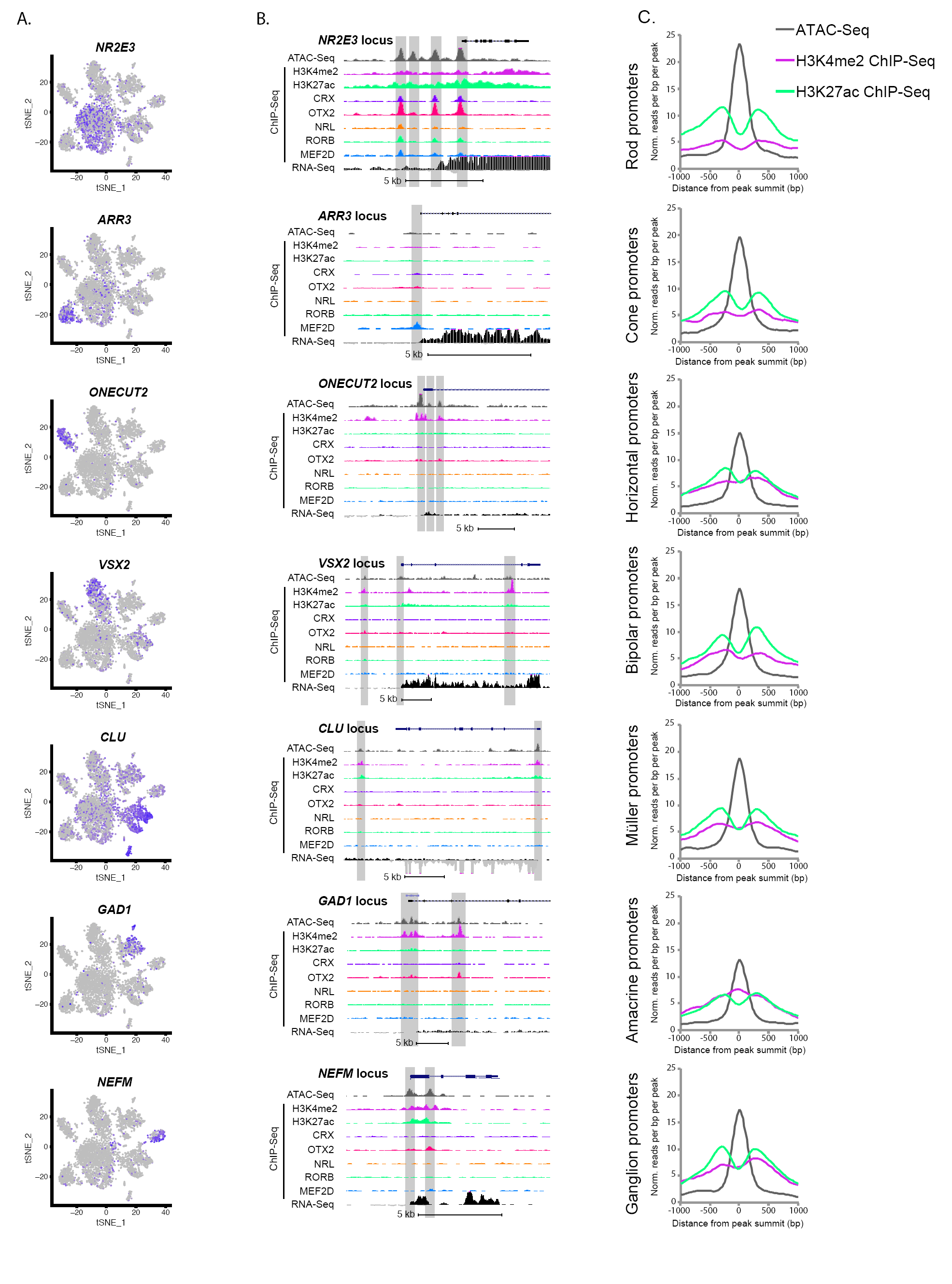
Related to Figure 4. **(A)** Marker gene expression of individual retinal cell types (*NR2E3*: rod photoreceptor cells; *ARR3*: cone photoreceptor cells; *ONECUT2*: horizontal cells; *VSX2*: bipolar cells; *CLU*: Müller glial cells; *GAD1*: GABAergic amacrine cells; *NEFM*: retinal ganglion cells). **(B)** Candidate regulatory elements at cell-type-specific gene loci. **(C)** Aggregate plots of ATAC-Seq or ChIP-Seq for H3K4me2 and H3K27ac at cell-type-enriched gene promoters.

**Figure S5.**
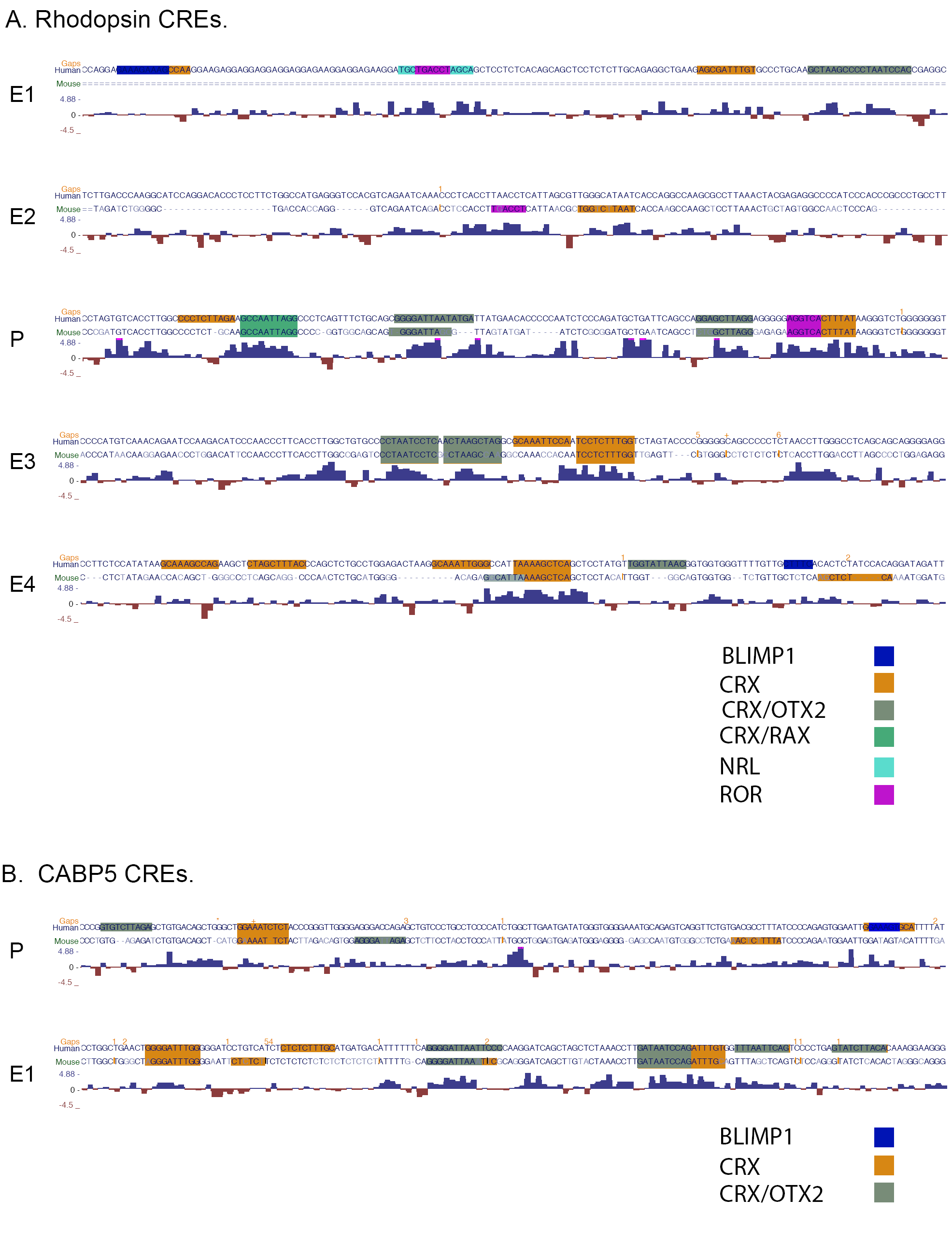
Related to Figure 5. **(A)** Human-Mouse genome alignment and conservation at candidate human *Rhodopsin* cis-regulatory elements. **(B)** Human-Mouse genome alignment and conservation at candidate human *CABP5 cis*-regulatory elements.

**Figure S6.**
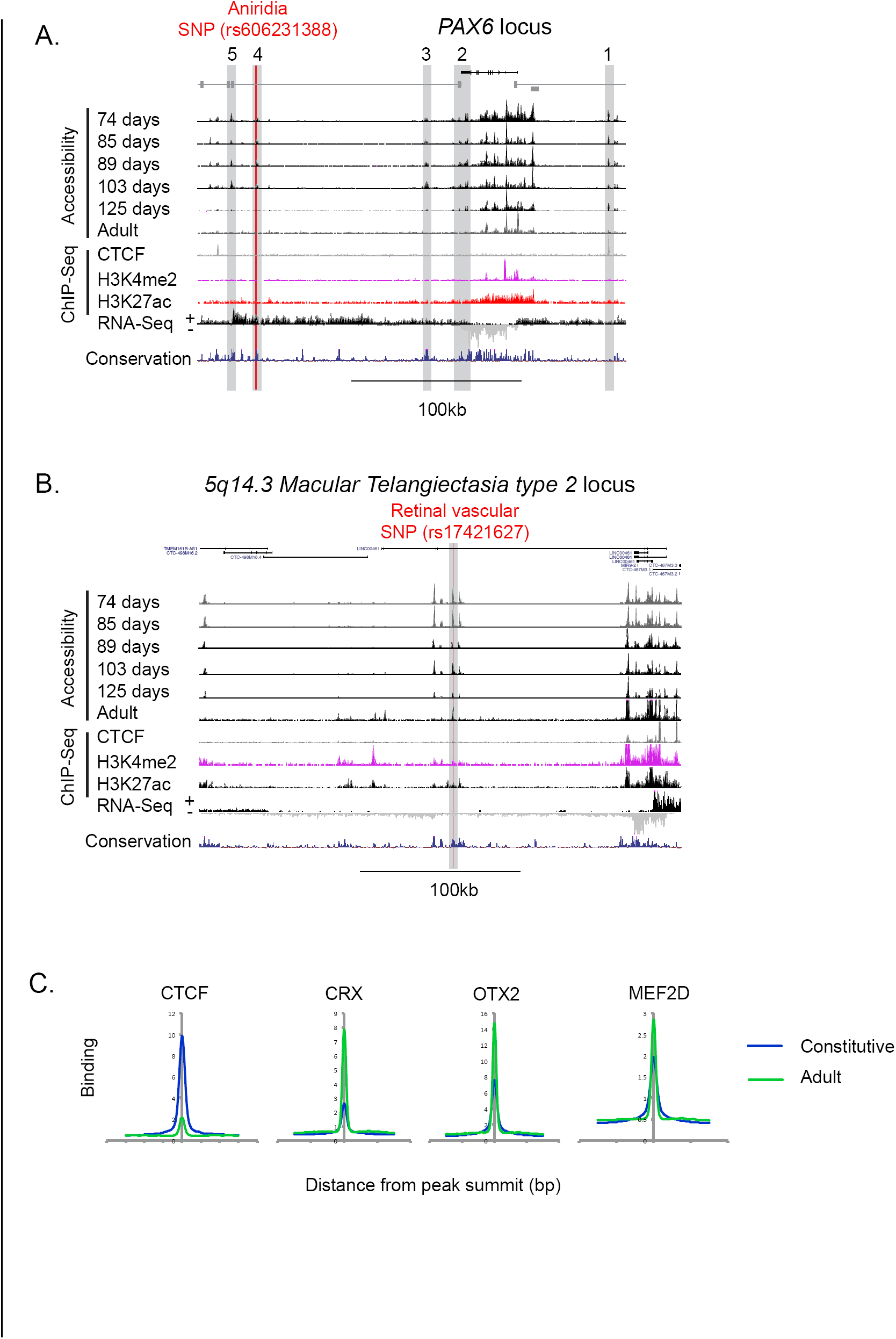
Related to Figure 6. **(A&B)** Developmentally dynamic DNA accessibility at the human *PAX6* locus or Macular Telangiectasia type 2 5q14.3 GWAS locus displayed as custom UCSC browser tracks for DNase-Seq (developing) or ATAC-Seq (adult) from a time course of human retinal development and CTCF, H3K4me2 and H3K27ac ChIP-Seq and total RNA Nuc-Seq from adult human retina (Red Line: SNPs resulting in aniridia and foveal hypoplasia (Bhatia et al., 2013) and associated with vascular anomalies and macular telangiectasia type 2 (Scerri et al., 2017). **(C)** Comparison of TF binding at constitutively or adult-only accessible DNA regions based on ChIP-Seq.

**Figure S7.**
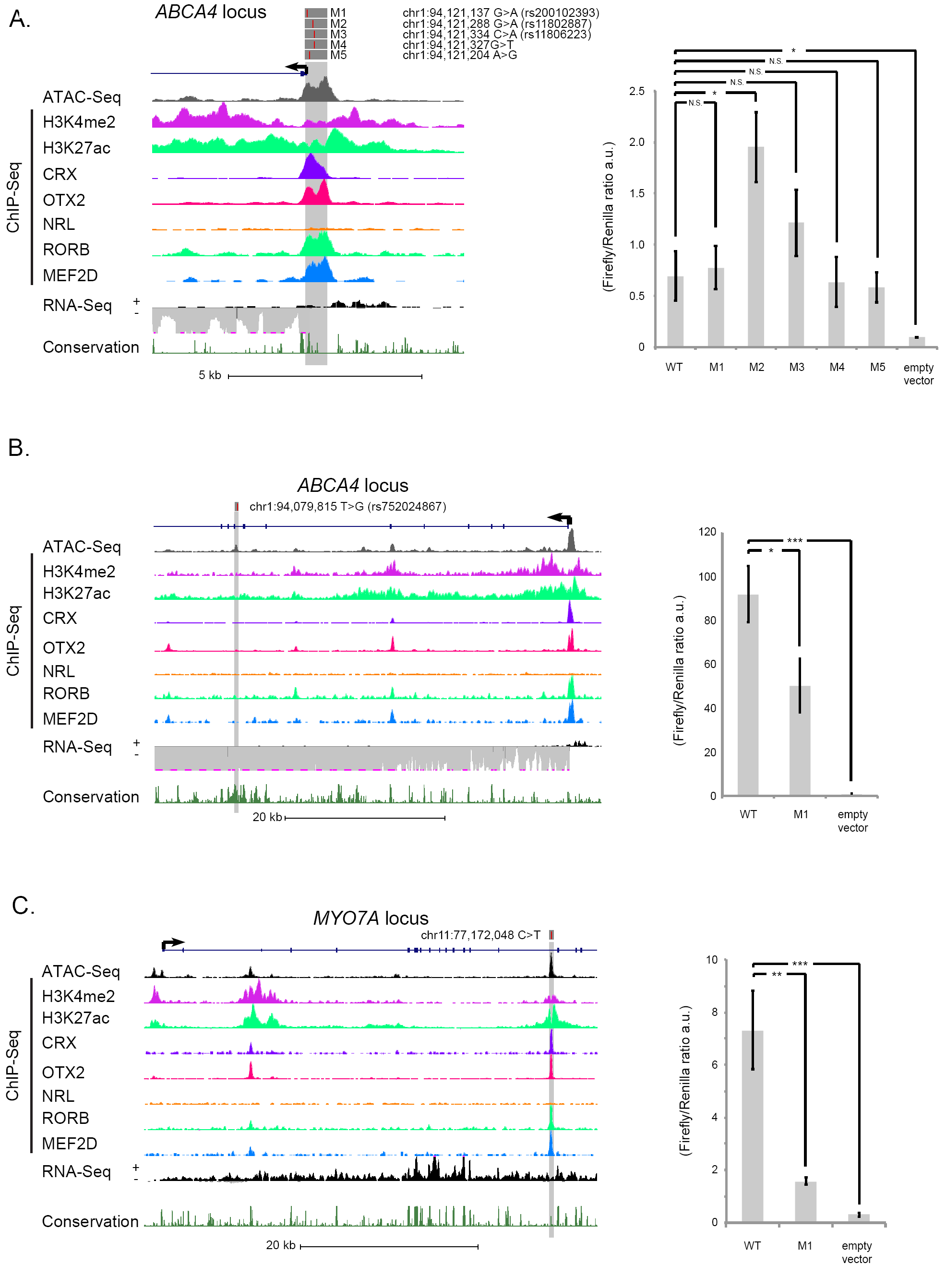
Related to Figure (A-C) *ABCA4* and *MYO7A* gene loci showing custom UCSC browser tracks for ATAC-Seq, H3K4me2, H3K27ac and TF ChIP-Seq and total RNA Nuc-Seq from adult human retina and conservation across 7 vertebrate genomes. Identified CREs highlighted in gray. Luciferase reporter assays comparing the normalized relative activities of ABCA4 consensus and variant promoters or enhancers or consensus versus variant MYO7A enhancer reporter constructs (*** p<0.001, * p<0.05, n=4). Genome coordinates according to build hg38.

